# Comprehensive benchmark of differential transcript usage analysis for static and dynamic conditions

**DOI:** 10.1101/2024.01.14.575548

**Authors:** Chit Tong Lio, Tolga Düz, Markus Hoffmann, Lina-Liv Willruth, Jan Baumbach, Markus List, Olga Tsoy

## Abstract

RNA sequencing offers unique insights into transcriptome diversity, and a plethora of tools have been developed to analyze alternative splicing. One important task is to detect changes in the relative transcript abundance in differential transcript usage (DTU) analysis. The choice of the right analysis tool is non-trivial and depends on experimental factors such as the availability of single- or paired-end and bulk or single-cell data. To help users select the most promising tool for their task, we performed a comprehensive benchmark of DTU detection tools. We cover a wide array of experimental settings, using simulated bulk and single-cell RNA-seq data as well as real transcriptomics datasets, including time-series data. Our results suggest that DEXSeq, edgeR, and LimmaDS are better choices for paired-end data, while DSGseq and DEXSeq can be used for single-end data. In single-cell simulation settings, we showed that satuRn performs better than DTUrtle. In addition, we showed that Spycone is optimal for time series DTU/IS analysis based on the evidence provided using GO terms enrichment analysis.

## Introduction

In higher eukaryotes, alternative splicing (AS) is an important process contributing to protein diversity. Different splicing events include exon skipping, alternative 3’/5’ splice site usage, mutually exclusive exon usage and intron retention. When a gene is spliced differently between two conditions, the relative abundance of a transcript can shift, irrespective of a change in the overall expression of a gene. In differential transcript usage (DTU), the distribution of transcript abundance changes, irrespective of a change in gene expression. DTU can have various functional consequences, e.g., switching from a protein-coding transcript to a non-coding transcript or switching between transcripts with different functions. Isoform switching is a special case of DTU, where we focus on DTU of the most abundant transcript [1].

DTU has been studied in various diseases. For example, Vitting-Seerup and Sandelin et al. showed that 19% of genes with multiple transcripts involve functional isoform switches in cancer [1]. Parkinson disease-related gene candidates were implicated in DTU, but these were not detected as differentially expressed genes, highlighting the importance of this type of analysis [2]. Since many tools have been proposed for DTU analysis, a key question is which tool to choose for a particular analysis.

To address this question, previous benchmark analyses have compared different workflows for DTU detection. Focusing on plant systems, Liu et al. compared differential splicing detection tools in simulated and real datasets [3]. Differential splicing events were simulated based on the changes indicated by relative transcript abundance in each gene using the Flux simulator [4]. DEXSeq and DSGSeq performed well with simulated data with area under the ROC curve (AUC) around 0.8 [5,6]. To evaluate the performance of tools in detecting novel, i.e. not previously annotated, events, the authors also performed DTU analysis with a truncated annotation. Cufflinks performed best with acceptable precision (0.9) and recall (0.7) on the *de novo* splicing events discovery. DICAST, a docker-integrated alternative splicing benchmark tool, allows users to compare splicing-aware mapping tools and splicing event detection tools on simulated and real data sets [7,8]. Similarly, a large-scale study by Jiang et al. focused on event-based tools applied to simulated datasets [9]. However, only tools that detect and quantify splicing events in one condition are included in the pipeline. In Merino et al., differential splicing tools were tested systematically in scenarios with differential splicing and/or differential transcript expression [10].

The authors concluded that DEXSeq [6] and LimmaDS [11] are the best tools for detecting DTU. However, the pipeline used the outdated tool TopHat [12] as aligner, whereas STAR [13] has been shown to perform better [14,15]. In a method paper by Love, Soneson and Patro, DEXSeq [6] and DRIMSeq [16] are used to perform DTU analysis [17]. These two papers only included five DTU tools, while we could currently find twelve tools for detecting DTU. The recent addition of new contenders motivated us to perform a comprehensive benchmark covering various experimental settings. In particular, we acknowledge a growing interest in single-cell DTU analysis, which has thus far not been covered in benchmarking analyses.

We further consider the challenging scenario of time series AS analysis, which was not previously covered in benchmark studies despite the importance of such analysis in recapitulating AS changes during development or in response to environmental changes. For example, time-dependent AS genes were detected in plants after exposure to cold temperatures, suggesting changes in night-to-day conversion and circadian control [18].

We compared twelve DTU detection tools, six of which had not previously been benchmarked. We utilized both simulated datasets and actual human transcriptomic datasets for this comparison. Our simulations covered various settings: in bulk settings, sequencing technology types (either single-end or paired-end), number of replicates (four or eight), and three background levels are considered. The term ‘background’’ refers to the likelihood of a gene not exhibiting differentially expressed transcripts, with a higher probability indicating a greater number of genes without DTU events. Our primary focus in the results section is on paired-end data. In single-cell settings, the number of cells and two background levels are considered. While we anticipate that paired-end sequencing excels in transcript detection, some studies still employ cheaper single-end sequencing, e.g. in time-series analysis, to support a larger sample number [19]. Understanding the performance of single-end data in DTU analysis is therefore crucial. In each bulk scenario, we simulated three scenarios contributing to transcript changes. We further categorized the results based on smaller or larger fold changes and the number of isoforms in a gene to understand the impact of different features on DTU detection. We simulated single-cell datasets to evaluate single-cell DTU tools such as DTUrtle and satuRn [20,21]. Additionally, we explored the qualitative differences between static pairwise comparisons and time series DTU analyses.

For paired-end sequencing, edgeR, DEXSeq, and LimmaDS emerged as top-performing tools [6,11,22]. In the context of single-end sequencing, we recommend DEXSeq and DSGseq [5]. LimmaDS was robust in detecting different types of DTU events (Figure 1). For time series data, Spycone was particularly effective in identifying biologically relevant events throughout the progression of a SARS-Cov-2 infected cell line. In single-cell data, satuRn has a better performance than DTUrtle. Taken together, this analysis provides a comprehensive view of the current state of DTU analysis in various scenarios.

**Figure 1.**
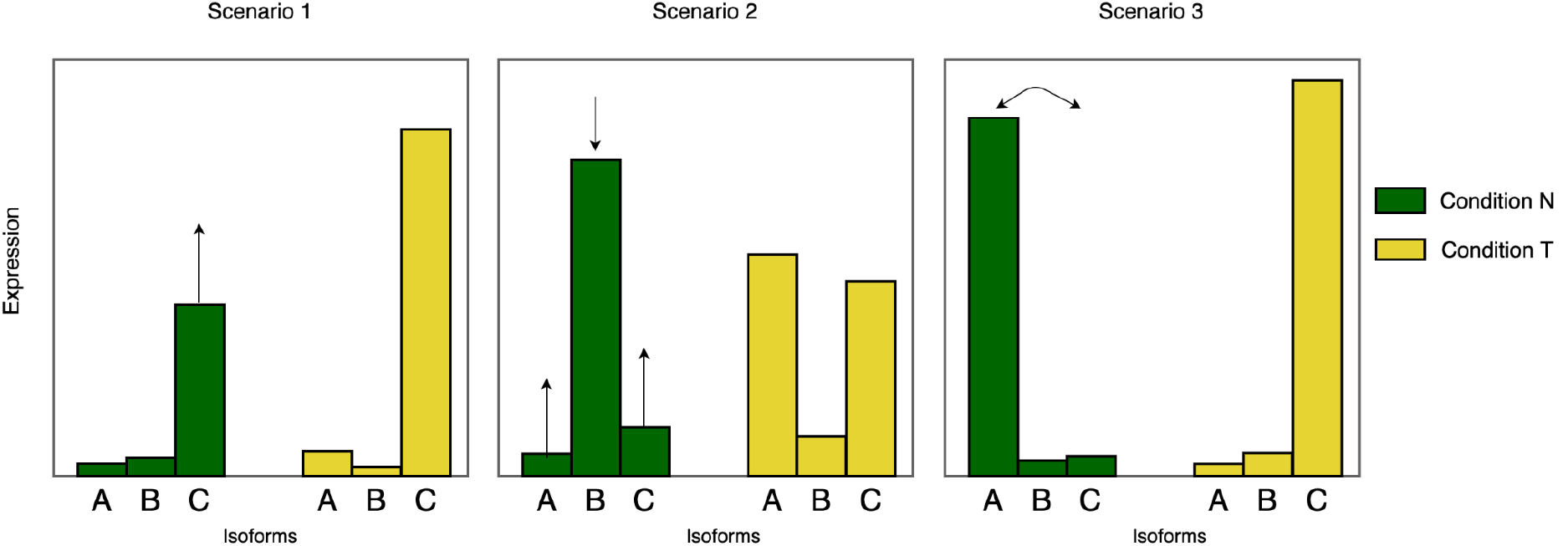
Scenario 1 (S1) are events that have one differentially expressed transcript, which are usually detected as differentially expressed genes as well. S2 are events that change the distribution of the transcript abundance between conditions. This change can involve multiple transcripts. S3 are events when the abundance is redistributed between two transcripts, it is called isoform switch (IS). The arrows indicate the changes represented in each scenario.

## Methods

### Simulation

Our simulation process is similar to the one proposed by Merino et al., i.e. we used RSEM (v1.3.3) to simulate single-end and paired-end data using estimated abundances that are inferred from sequencing model parameters from real datasets and reference transcriptome [10]. The rsem-calculate-expression function estimates model parameters from real datasets [23]. The function collects statistics from the dataset, including the number of reads, the number of reads aligned to multiple and unique loci, the read and fragment length distribution and the quality score distribution.

The single-end data model parameters are estimated from GSE157490 [19], a cell line dataset with SARS-Cov2 infection sequenced at 100M reads. The paired-end data used to learn parameters is GSE162562 and GSE190680, which is also a dataset of patients with SARS-Cov2 infection sequenced at 100M reads [24,25]. Each dataset is simulated with 50 million reads, which is the minimum depth to robustly detect DTU [26] and 100 million reads.

Baseline transcript expression levels are taken from the SARS-CoV2 datasets. Next, we adjust the transcript counts to generate simulated data for three DTU scenarios (Figure 1). Since changes in transcript expression and DTU are confounded by changes in the overall expression of a gene, we consider both effects together. First, for each gene, we consider a random fold change ranging from 2 to 5 between conditions. The transcript ratios are generated using a Dirichlet distribution, which describes the probabilities of k categories given a density distribution with k dimensions. This approach is ideal for simulating transcript ratios as the sum of the vectors is 1. k represents the number of transcripts in a gene, with each transcript being assigned an expression value. The higher the probability associated with transcript i, the higher the expression value. To have a higher statistical power for detecting DTU transcripts, we simulate DTU transcripts with higher expression level. The following formula shows the simulation of expression value for each transcript i in condition j.

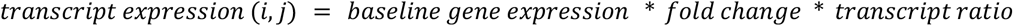

Note that for the baseline (e.g. a control), the fold change is 1, whereas for the condition of interest we consider the random fold change. For creating replicates with measurement noise, we compute the expression values using a negative binomial distribution, where the dispersion for each transcript is estimated using DESeq2

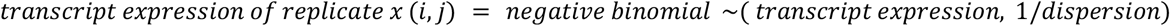

We split the genes across scenarios to obtain a mixture for the final data set. In theory, we could consider data sets that only consider individual scenarios, but this would not result in a realistic data set for evaluating the tools. In the next step, we must modify the transcript ratios according to the scenarios we consider. For scenario S1, only a single transcript of the gene changes expression. For scenario S2, more than two transcripts are subject to changes in relative abundance. In scenario S3, the relative abundance of two transcripts is swapped, signifying an isoform switch event.

We considered three background levels with an increasing fraction of genes whose expression remains unchanged: 0, 0.1, and 0.5. The modified transcript results are then used for the simulation. The rsem-simulate-reads command is used for the simulation. RSEM reference is generated with the human genome GRCh38, theta0 parameter, noise proportion to the background is set to 0.1.

In our study, we simulated a total of four conditions, incorporating various parameter combinations (Table 1). 100M reads are simulated only with four replicates and background level 0.5.

**Table 1.** Different simulated data generated

| Type | replicates | depth | background |
| --- | --- | --- | --- |
| paired-end | 4 | 50M, 100M | 0, 0.1, 0.5 |
| paired-end | 8 | 50M | 0, 0.1, 0.5 |
| single-end | 4 | 50M, 100M | 0, 0.1, 0.5 |
| single-end | 8 | 50M | 0, 0.1, 0.5 |

### Single cell simulation

We used the same simulation workflow for a single cell, except the model parameters for RSEM are learned from a demultiplexed Smart-seq2 dataset derived from human cells [27]. We adapted our simulation method for single-cell transcript counts. In particular, we used the *–single-cell-prior* parameters in RSEM to model sparse matrices. RSEM seems to perform better in simulating Smart-seq2 dataset (Figure S2). To test the methods with simple single-cell data, we used two cell types with a population of 900 to simulate. We simulated balanced datasets with 20, 50, 100, 200, 500 and 1000 cells containing the same amount of cells for each cell type. We simulated DTU events in single-cell with two background levels - 0 and 0.1. Background level here also means the percentage of expressed genes that will stay unchanged between the two cell types.

### Differential transcript usage methods

We performed an in-depth literature search in databases PubMed and Google Scholar with keywords such as “differential transcript usage”, “differential isoform usage”, “isoform switch”. We selected publications that describe a novel method for DTU analysis for bulk transcriptomics data. Iso-DOT is excluded due to long runtime (>20 days without parallelization) [28], rSeqDiff is excluded due to not supporting replicates [29] and IUTA is excluded due to incompatibility with STAR output [30].

Tools that can detect DTU can be categorised into exon/junction-centric (JunctionSeq [31], seqGSEA [32], DSGSeq [5]) and transcript-centric (DEXSeq [6], DRIM-Seq [16], DTUrtle [20], iso-KTSP [33],satuRn [21], NBSplice [34], edgeR [22], LimmaDS [11]), as well as assembly-based (Cufflinks/cuffdiff[35]).

Exon-centric tools like DEXSeq (v1.24.0) and JunctionSeq (v1.5.4) use a generalized linear model to analyze differences in exon and splice junction usage. DEXSeq, originally designed for exon counts, also works with transcript counts. It uses a formula to compare conditions and identifies significant genes based on adjusted p-values. JunctionSeq also uses adjusted p-values for gene evaluation. DSGSeq (v0.1.0) compares exon counts between conditions using negative binomial statistics. At the same time, seqGSEA (v1.36.0) combines DSGseq and DESeq methods, using exon counts and a rank-based strategy to output p-values for differential transcript usage (DTU) genes. Transcript-centric tools include DRIMSeq (v1.24.), which uses a dirichlet-multinomial model for transcript abundance analysis, and DTUrtle (v1.0.2), which adds extra filtering steps and uses StageR for more accurate gene-level false discovery rate correction. Iso-KTSP (v1.0.3) identifies transcript pairs that differentiate conditions, scoring them based on expression frequency. satuRn (v1.4.2) models transcript counts using a quasi-binomial model, and Cufflinks/cuffdiff (v2.2.1) aligns transcripts de novo and analyzes differential expression. For gene significance, tools using adjusted p-values consider values below 0.05 as significant. Iso-KTSP and DSGseq, which don’t provide p-values, use cutoffs of 0.5 and 5 based on author recommendations. These tools are listed in Table 2.

**Table 2.** All DTU tools published after 2010.

| Tool | Implementation | Year | Reference | Principle idea | Excluded |
| --- | --- | --- | --- | --- | --- |
| DEXSeq | R | 2012 | [6] | Negative binomial generalized linear model to model transcript counts |  |
| DRIMSeq | R | 2016 | [16] | Use dirichlet-multinomial model to model relative abundance of transcript |  |
| seqGSEA | R | 2014 | [32] | Negative binomial models in DSGSeq and DESeq |  |
| DTUrtle | R | 2021 | [20] | Dirichlet-multinomial model from DRIMSeq |  |
| Junction Seq | R | 2016 | [31] | Negative binomial generalized linear model to model junction counts |  |
| NBsplice | R | 2020 | [34] | Negative binomial generalized linear model to model transcript counts, and test with a linear hypothesis |  |
| satuRn | R | 2021 | [21] | Quasi-binomial generalized linear model to model transcript counts |  |
| limmaDS | R | 2013 | [11] | Apply linear model to detect transcript changes |  |
| Cuffdiff2 | C++ | 2012 | [35] | Use a poisson model to estimate changes in transcript counts |  |
| iso-KTS P | Java | 2014 | [33] | Classify transcripts to condition specific based on the change of transcript abundance |  |
| DSGSeq | R | 2013 | [5] | Negative binomial statistics to detect transcript changes |  |
| edgeR | R | 2010 | [22] | Negative binomial generalized linear model to model transcript counts |  |
| IUTA | R | 2014 | [30] | Estimate transcript usage, followed by testing DTU under Aitchison geometry | Incompatible with STAR |
| IsoDOT | R | 2015 | [28] | Estimate transcript usage with a penalized regression method | Long run time |
| rSeqDiff | R | 2013 | [29] | Apply linear poisson model to estimate transcript counts | Not supporting replicates |
| Time series tools |  |  |  |  |  |
| TSIS | R | 2017 | [36] | detection and characterization of isoform switches for time series data |  |
| Spycone | python | 2023 | [37] | detection and characterization of isoform switches for time series data |  |

### Differential transcript usage methods for time series data

There are two time series tools for detection of isoform switches in time series data: TSIS [36] and Sypcone [37]. TSIS and Spycone are the tools that detect switch points between transcript pairs and calculate adjusted p-values based on the replicates. Then, it applies filters to select features like switching probabilities (i.e. the ratio of samples that has a higher relative abundance in one transcript than the other), the difference of transcript expression before and after the switch. In Spycone, additional metrics are calculated such as event importance and domain difference. To detect DTU genes in TSIS, the following filtering metrics are used by default: 1) probability of switching > 0.5, 2) difference of expression before and after switching > 1, 3) p-value < 0.05, 4) correlation coefficient > 0.5. For the usage of Spycone, DTU genes are filtered with default parameters: 1) difference of relative abundance before and after switch > 0.2, 2) adjusted p-value < 0.05, 3) dissimilar correlation > 0.5 and 4) event importance > 0.3. For DEXSeq, we used a log-likelihood test with a reduced model. For LimmaDS, each time point is treated as a factor in multiple conditions.We used clusterProfiler R package to perform GO term enrichment analysis [38].

### Preprocessing and quantification

STAR (v2.7.8a) is used to align the reads to the genome (GRCh38 v107). We used STAR, which is currently among the best tools for splice-aware alignment [7]. Salmon (v1.7.0), kallisto (v0.44.0), RSEM and Cufflinks (v2.2.1) are used for transcript quantification. HTSeq (v2.0.1) is used to quantify exon counts for the input of seqGSEA. The workflow is shown in Fig. 2.

**Figure 2.**
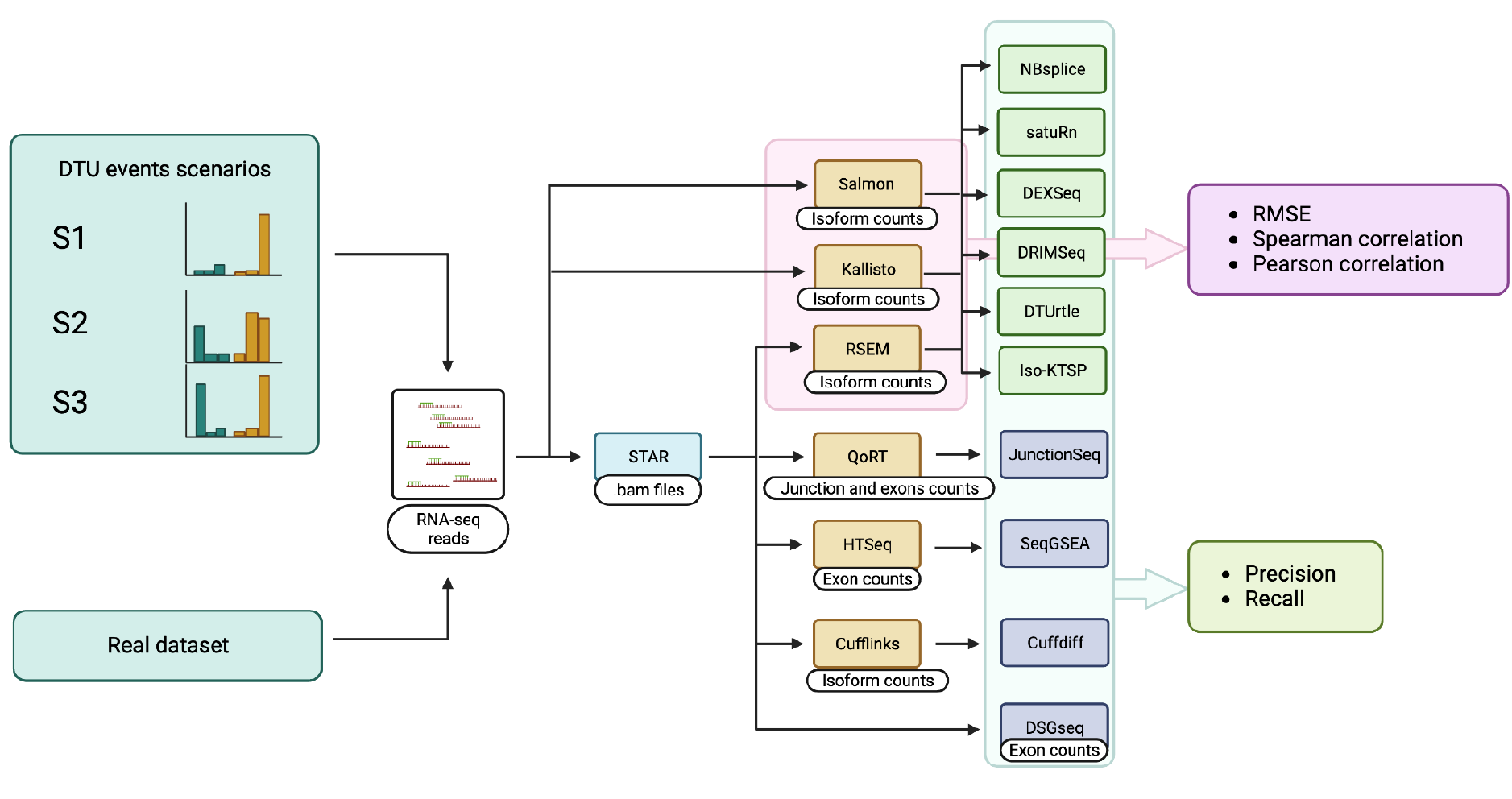
Analysis workflow of the methods. Different scenarios are simulated to evaluate the performance of the tools. STAR is used to map RNA-seq reads and the resulting bam files are used by RSEM, QoRT, HTSeq and cufflinks to generate transcript counts, junction and exons counts. Salmon and Kallisto directly use RNA-seq reads in fastq format to generate transcript counts. Transcript counts derived from Salmon, Kallisto and RSEM are compared by calculating root mean squared error (RMSE), Spearman and Pearson correlation. We generate DTU detection results from NBsplice, satuRn, DEXSeq, DRIMSeq, DTUrtle, and Iso-KTSP using transcript counts from Salmon, Kallisto and RSEM. Other DTU tools use the corresponding count tables. All DTU detection results are compared by calculating precision and recall compared to the simulated ground truth.

### Evaluation of performance

To evaluate the effectiveness of the tools on simulated data, we employed precision and recall as our performance metrics. We obtained a list of genes identified as having DTU, based on an adjusted p-value of less than 0.05, or surpassing a specific threshold in the case of iso-KTSP and DSGseq. These genes are positive. True positives (TP) are those among the positive genes that also appear in the simulated ground truth, while the rest are categorized as false positives (FP). Conversely, genes that are simulated with DTU but remain undetected by the tools are labeled as false negatives (FN). For stratification analysis, each event scenario is calculated separately. For different DTU events scenarios, P are genes simulated with the corresponding scenario. The formulas for calculating precision and recall are as follows:

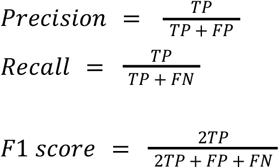

To evaluate the performance in a real dataset, a prostate tumor dataset (GSE222260) is used. The dataset consists of 10 normal tissues and 20 prostate carcinoma tissues.

## Data availability

The RNA-seq data of prostate tumor dataset is available from GSE222260 [39] and data from patients with SARS-Cov-2 infection is available from GSE162562 and GSE190680 [24,25]. Both datasets were preprocessed and analyzed as mentioned above. The time series RNA-seq data of SARS-Cov-2 infection data was obtained from GSE157490 [19]. The data is processed as described in [40].

## Code availability

Code for simulation, analysis and plot generation are available at https://github.com/yollct/diffIsoUsage_benchmark under the terms of the GNU General Public License, Version 3.

## Results

### Overall performance of DTU detection

In this study, we evaluated twelve DTU tools using simulated datasets that incorporated various scenarios, including single-end or paired-end data, four or eight replicates, and three distinct background levels.

The datasets included three DTU scenarios for differentially expressed genes. We assessed precision, recall and F1 scores for each scenario based on the significant results obtained, as detailed in Table 1. For tools providing adjusted p-values, a threshold of 0.05 was used to identify positive results, while for iso-KTSP and DSGseq, the thresholds were set at 0.8 and 5, respectively. The transcript counts from each simulated dataset were compared against a ground truth outlined in the supplementary file (Figure S1).

Figure 3 presents the metrics for the DTU detection tools on simulated data with four and eight replicates. Generally, an increase in recall was observed with eight replicates. DRIMSeq, DTUrtle, and JunctionSeq demonstrated comparable performances. NBSplice showed the highest precision overall, even with a background of 0.5 in four replicates, though its recall was low. LimmaDS achieved the highest recall (>0.2) in both sets of replicates. However, iso-KTSP’s precision decreased to 0.4 as background increased. In contrast, DEXSeq and satuRn showed less impact on precision at a background of 0.5 (Figure 3A). When evaluating with F1 scores, iso-KTSP appears to be the best performing tool despite low recall (Figure 3B).

**Figure 3.**
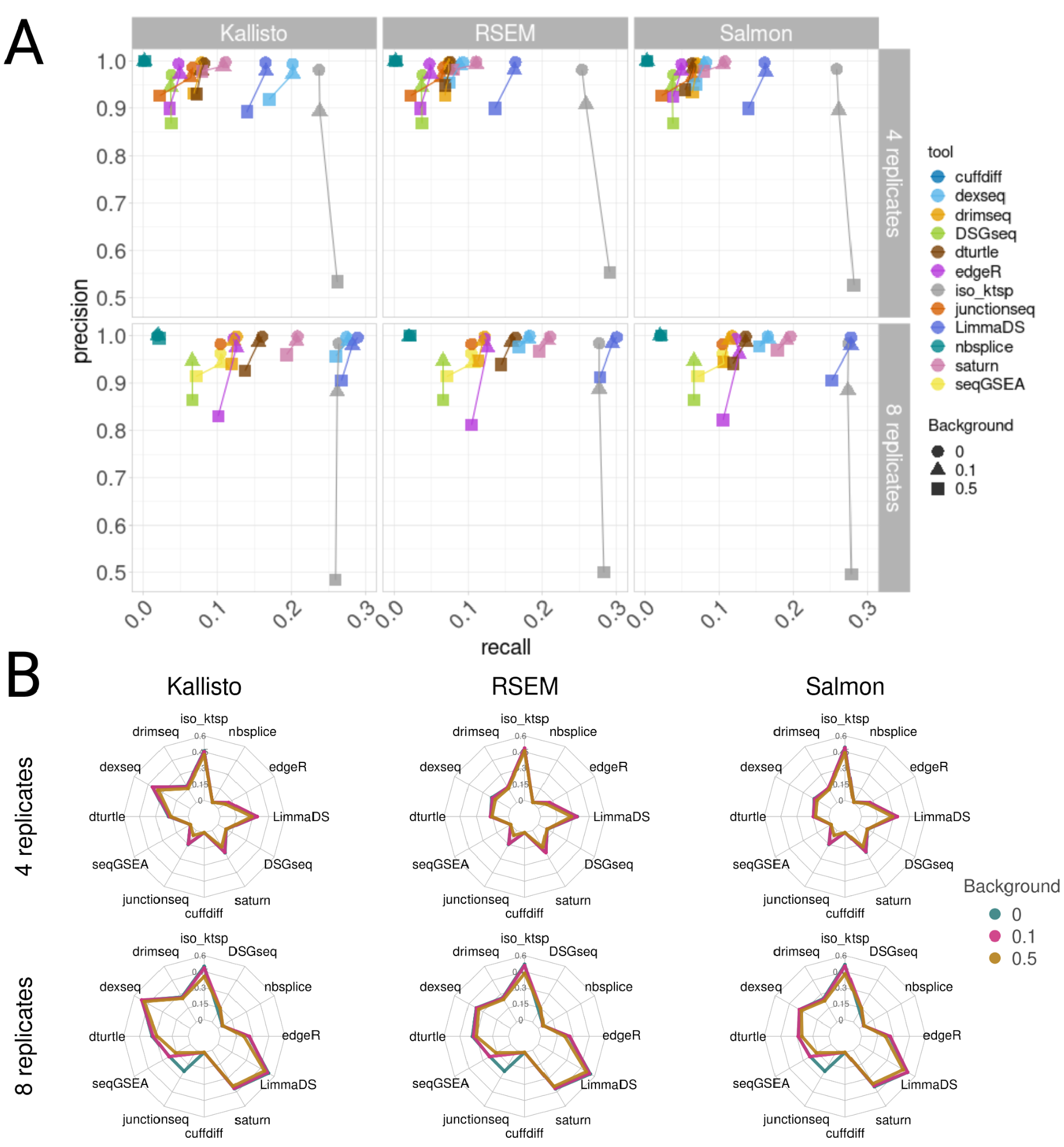
Metrics plot of all combinations of quantification tools and DTU methods from paired-end data with 4 replicates (upper-row) and 8 replicates (lower-row). A) Precision and recall plot. B) Radar plots showing the corresponding F1 scores.

Figure S4 illustrates the results for single-end data, where all tools exhibited low recall in four replicates, possibly due to the lack of additional information from the paired-end protocol. This limitation can be mitigated with more replicates (eight). edgeR and DEXSeq achieved the best precision in four and eight replicates, respectively. Notably, DEXSeq combined with Kallisto showed promising results in both sequence types (Figure S5). Similar to paired-end data, DEXSeq with

Kallisto excelled in both four and eight replicates. However, LimmaDS displayed low recall in all cases (close to 0), and its precision significantly dropped to 0.4 when the background was set at 0.5 (Figure S12).

To investigate the effect of increasing sequencing depth, we simulated 100M reads for 4 replicates at 0.1 background. Most tools have increased recall. LimmaDS has the most improvement (+0.1), but precision dropped. DSGseq has increased precision and decreased recall (Figure S19).

### Performance of tools in different stratifications

Figure 4 presents the F1 scores for various event types based on data quantified by Salmon. The results from Kallisto and RSEM quantifications are provided in the supplementary figures (Figures S6-8). The ranking of tools according to F1 scores are the same, except DEXSeq with kallisto quantification has the best performance. Additionally, we have stratified the genes according to the number of transcripts (Figure S3-5) and the extent of the fold change (Figure S9-11). F1 scores are determined using the ground truth for each specific category. While iso-KTSP demonstrates improved performance in Figure 4, this is not reflected in Figure 3A, where its tendency towards lower recall needs to be taken into account.

**Figure 4.**
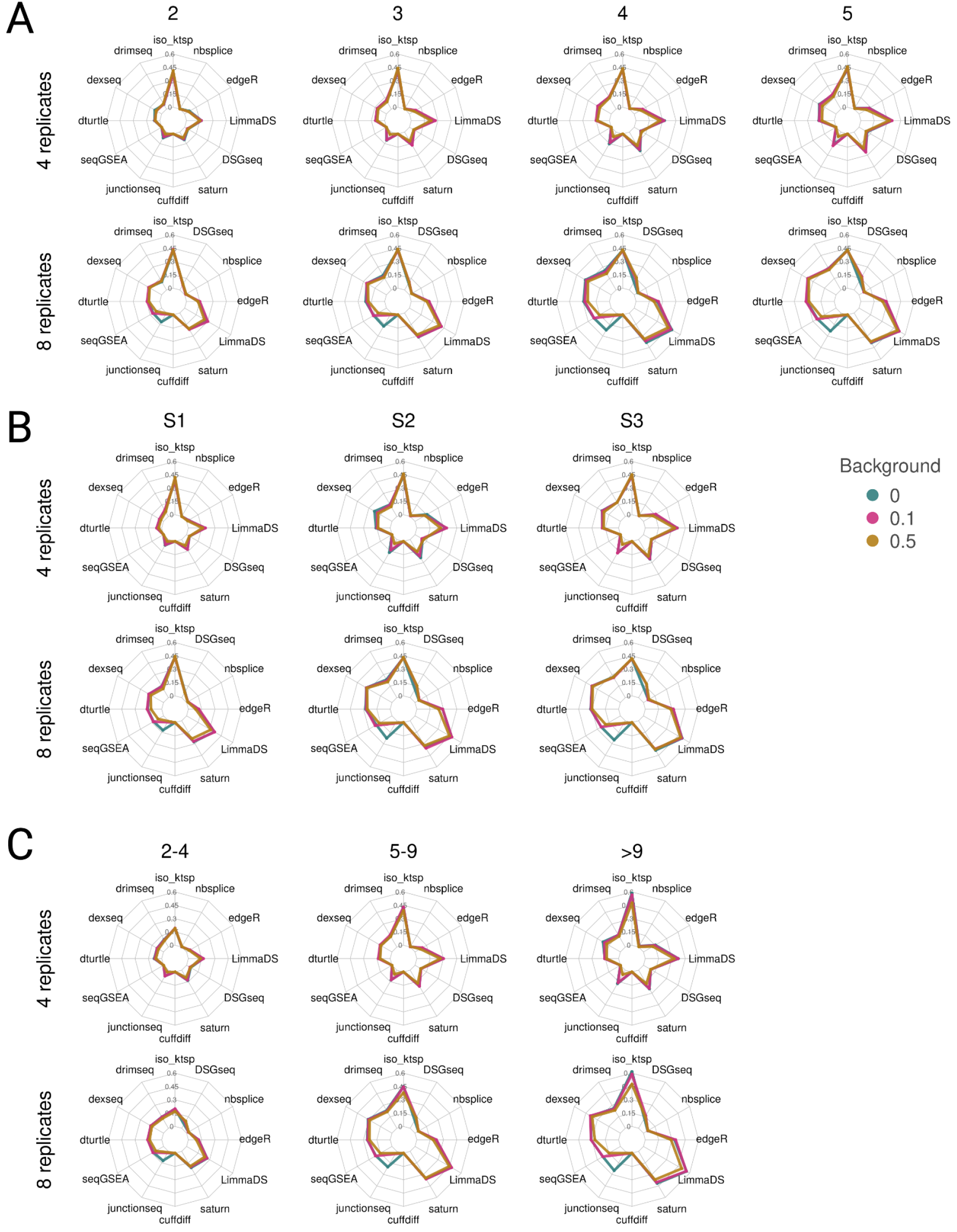
Radar plots showing F1 scores calculated after stratification for all combinations of quantification tools and DTU methods in paired-end data with 4 replicates (top row) and 8 replicates (bottom row). This result is derived from Salmon quantification. A) DTU events stratified by different scenarios B) DTU events stratified by fold change. C) DTU events stratified by number of transcripts.

As expected, the performance of all tools improves with eight replicates. LimmaDS, in particular, achieves the highest F1 score and recall (>0.4) for S2 and S3 DTU events (Figure S9, 11). When eight replicates are used, satuRn maintains high precision, even with a high background (Figure S10). When paired with Kallisto, DEXSeq outperforms LimmaDS. Generally, an increase in fold change magnitude correlates with higher detection of events, thereby boosting the F1 scores. Nonetheless, iso-KTSP’s F1 score remains consistent regardless of fold change. Moreover, satuRn demonstrates superior precision. Subsequently, we categorized genes into groups based on the number of transcripts, revealing a direct correlation between the number of transcripts and F1 scores. However, this categorization appears to have minimal impact on precision (Figure S4).ev In the analysis of single-end data, DEXSeq combined with Kallisto consistently exhibited superior performance in both four and eight replicate scenarios, as shown in Figure S13. Specifically, DEXSeq achieved an F1 score of 0.2 for S1 DTU events and approached 0.6 with eight replicates. iso-KTSP maintained steady F1 scores across all cases, similar to its performance in paired-end data, but it also identified false positives, as indicated in Figure S12. In contrast, LimmaDS showed weaker results in single-end data. For quantifications using RSEM and Salmon, seqGSEA outperformed others with an F1 score of 0.2 in eight replicates, while JunctionSeq led in the four-replicate category. Similar to paired-end data, a positive correlation between fold change and F1 scores was observed (Figure S14). However, the number of isoforms appeared to have a minimal impact on the results (Figure S15).

### Benchmarking with a real transcriptome dataset

In our study, we utilized the paired-end prostate cancer dataset, previously used by Merino et al., to evaluate various DTU tools. We quantified sequencing reads using Salmon, Kallisto, and RSEM,applying all DTU tools except JunctionSeq, which was excluded due to its excessive runtime. Figure 5 illustrates the overlap in DTU gene sets detected by different tools, along with the F1 scores derived from our simulation analysis (Figure 5).

**Figure 5.**
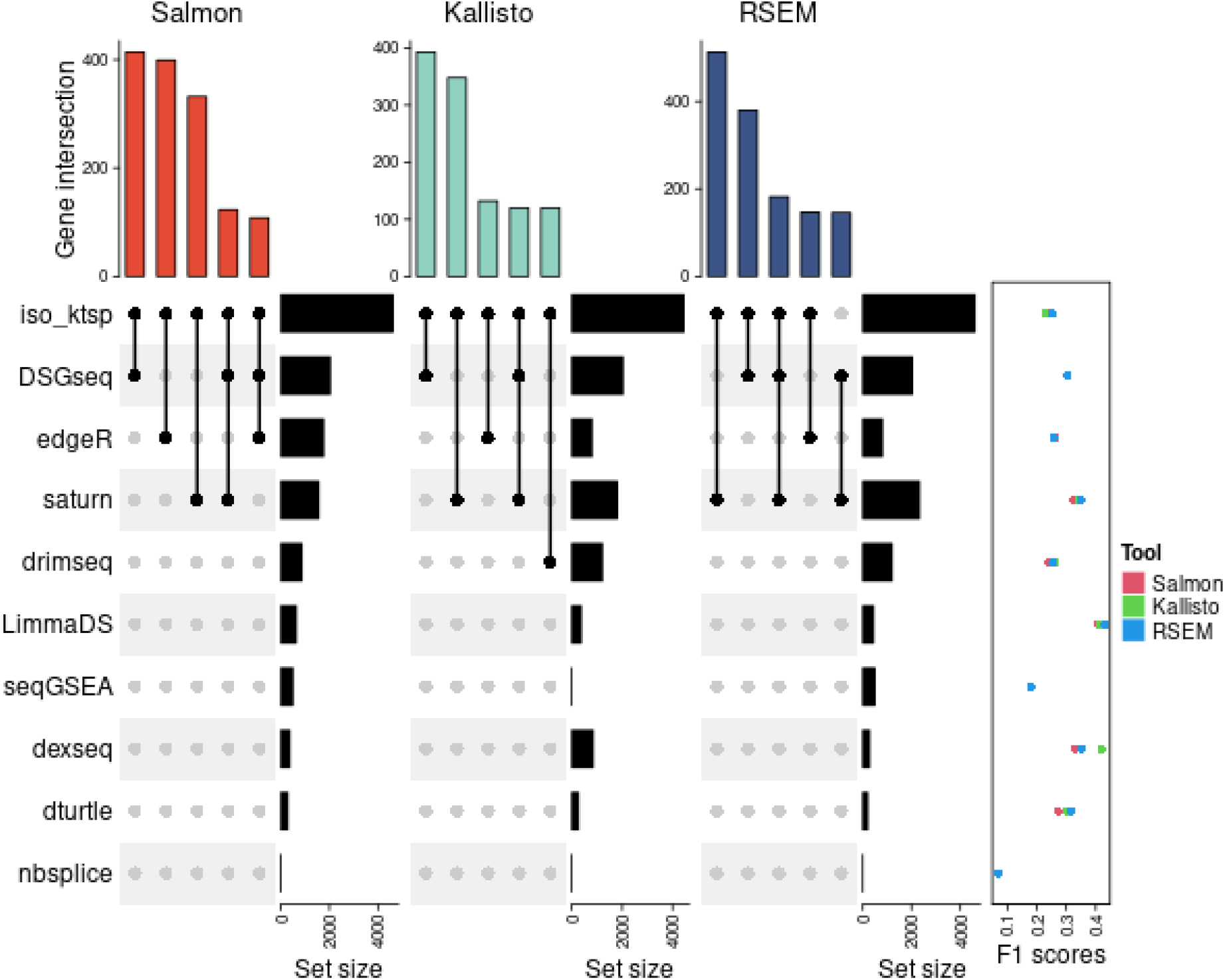
Upset plots showing the overlapping DTU genes found by each tool Cancer dataset obtained from [39] are analyzed with the workflows. Each upset plot shows the number of DTU genes detected for each tool and the overlaps among them. The right dot plot shows the F1 scores obtained from the simulated dataset, with 8 replicates and a background of 0.5.

Iso-KTSP identified the highest number of DTU genes, but the F1 score was relatively low (0.25), indicating a higher rate of false positives in the simulated data. Conversely, LimmaDS, with a higher F1 score (0.4), detected only a limited number of genes in this dataset. We observed variations in the number of genes detected by different DTU tools depending on the quantification method used. For instance, DEXSeq identified the most genes with Kallisto counts, while edgeR and LimmaDS detected the most with Salmon counts, and satuRn found the most with RSEM counts. Notably, NBSplice did not detect any significant genes in Salmon and Kallisto, and only one gene in RSEM.

### Performance in single-cell data

We assessed the performance of DTUrtle and satuRn on single-cell data using a simulated dataset created with the specified method. Each dataset comprised two cell types, each with an equal number of cells. We calculated precision and recall based on the simulated ground truth. Figure 6 illustrates the precision and recall as the number of cells in each cell type increases. Generally, we note a high precision (around 0.9) when there are 50 or more cells in each cell type. The recall, however, shows agradual increase with the rising number of cells. SatuRn demonstrated a higher recall than DTUrtle, reaching a recall of 0.9 when each cell type had 500 cells, compared to DTUrtle’s 0.73. As the background level rises, both precision and recall decline. SatuRn registered a precision of 0.83 and a recall of 0.88, whereas DTUrtle posted a precision of 0.93 and a recall of 0.69. However, with a higher background level, an increase in the number of cells resulted in a slight dip in precision (a decrease of 0.06 from 100 to 200 cells and a further decrease of 0.01 thereafter). In addition, we performed a pseudo-bulk analysis where transcript counts are aggregated into meta cells. The meta cells are then analyzed using methods designed for bulk data. The result shows that this approach does not improve precision and recall (Figure S17).

**Figure 6.**
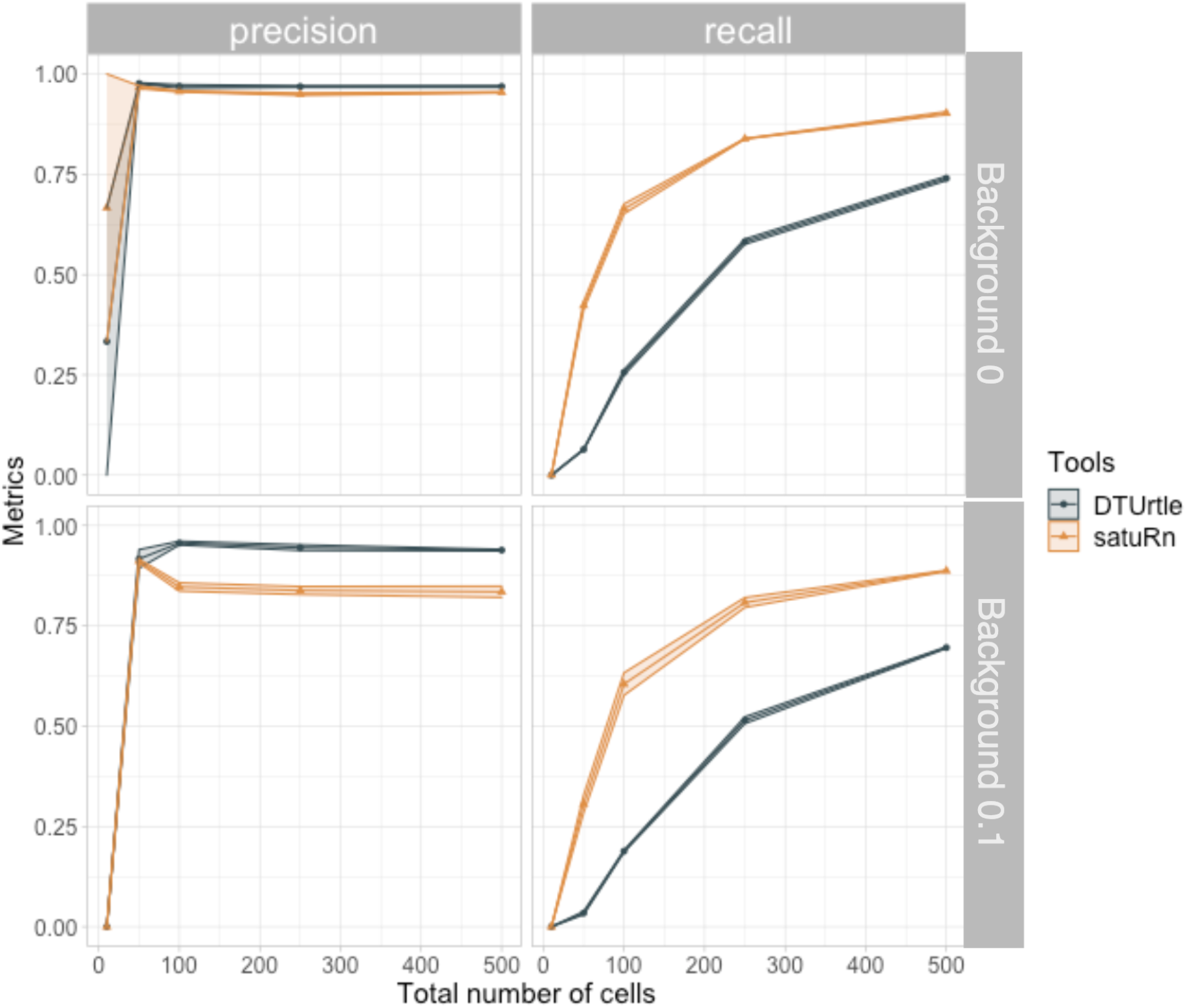
Precision and recall of satuRn and DTUrtle for single-cell simulation. Each plot consists of the result from a different number of cells (x-axis). Precision from simulation with background level 0 (top left). Recall from simulation with background level 0 (top right). Precision from simulation with background level 0 (bottom left). Recall from simulation with background level 0.1 (bottom right).

### Time series isoform switch analysis

Another method for inferring isoform switches is time series isoform switch detection. With time series data, we can extract dynamic changes in transcript usage. Here, we applied DEXSeq, LimmaDS, satuRn, TSIS and Spycone on time series single-end transcriptomic data for SARS-Cov-2 infection with eight time points and four replicates [19]. Figure 7 shows the results of the comparison. In all the comparisons, DEXSeq and LimmaDS have many overlapping DTU genes (1731). edgeR didn’t find any significant genes. Spycone and TSIS, which are specifically designed for time series data, have only a few overlaps (Figure. 7A). In GO terms biological processes enrichment of the significant DTU genes, the terms enriched in each of the sets are different, in which DEXSeq, LimmaDS and TSIS are enriched in generic cellular functions such as cadherin binding, ubiquitin-related terms etc (Figure. 7B). satuRn has similar but less enriched The Spycone gene set is enriched in MHC protein complex binding, which is essential to adaptive immunity. For the term IgA bindings, IgA are found to be as part of the early humoral immune response to neutralize SARS-Cov-2 virus upon infection [41].

**Figure 7.**
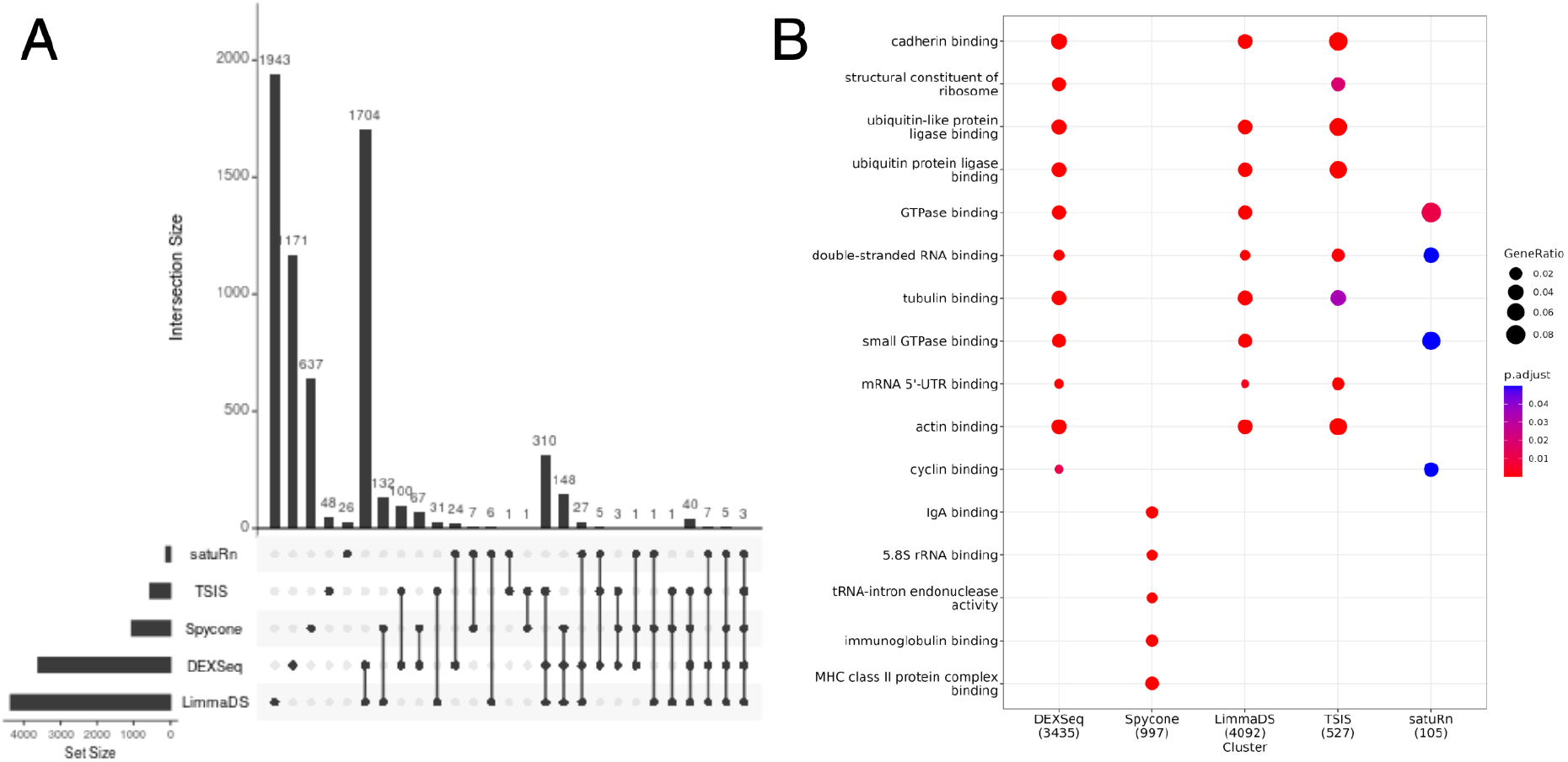
A) An upset plot showing the overlapping DTU genes detected by DEXSeq, TSIS, NBSplice, LimmaDS and Spycone time series isoform switch detection. B) Dotplot showing the enrichment terms from the DTU genes detected by all tools. NBSplice didn’t report significant genes.

## Discussion

In this analysis, we conducted a comprehensive benchmark study using both simulated data and real transcriptomics datasets, considering both static and dynamic conditions. Our simulation approach aimed to mimic real datasets, including the presence of stochastic noise. To achieve this, we simulated three background levels: 0, 0.1, and 0.5, representing the proportion of expressed genes that remain unchanged between two conditions. Additionally, we simulated both paired-end and single-end RNA-Seq data with four and eight replicates.

Our simulation approach contrasts with that of Merino et al. [10], where the authors specifically assigned abundance to each transcript between two conditions to simulate differential transcript abundance. Instead, we adopted a dirichlet-multinomial model to simulate abundance based on transcript probabilities. The Dirichlet-multinomial model effectively manages transcript expression as multivariate count data, accommodating overdispersion to more accurately reflect real-world data scenarios. Furthermore, while Merino et al. focused solely on simulating DTU events where two transcripts change in abundance and evaluated them using DTU tools, we simulated events with more than two transcripts change in abundance (S2) and assessed their performance with DTU tools.

In the simulated scenarios, we observed that single-end data exhibited lower recall (Figure S12). This decrease in recall can be attributed to the limitations of single-end sequencing, which cannot capture both ends of a cDNA fragment, leading to a reduced probability of observing junction reads. Consequently, single-end data has lower power for detecting the correct transcript. In our simulations, we found that single-end data could detect approximately 10,000 transcripts, while paired-end sequencing detected around 30,000 transcripts at a depth of 50M reads (Figure S18). However, despite its higher sensitivity, paired-end sequencing faces challenges in transcript detection due to transcriptional noise stemming from transcriptional stochasticity [42]. Furthermore, short reads cannot accurately detect transcript counts due to the increased likelihood of mapping reads to multiple genomic locations. This limitation can be mitigated by long-read sequencing technology once it becomes more prevalent [43].

Another factor influencing our observations is the simulation methods employed. RSEM served as the simulation engine, and since it is one of the quantification methods used in our analysis, there may be a bias in favor of RSEM. In our simulation approach, we used the mean expression values for each transcript from a real transcriptome dataset. Additionally, our simulation approach utilized a Dirichlet-multinomial model to generate transcript abundances, which might favor tools that also utilize the Dirichlet-multinomial model for modeling, such as DRIMseq and DTUrtle.

In our findings, we noted that tools employing a generalized linear model, such as NBSplice, DEXSeq, satuRn, JunctionSeq, and edgeR, as well as those utilizing a Dirichlet-multinomial model, such as DRIMSeq and DTUrtle, typically exhibited superior precision. Conversely, the only tool employing a linear model, LimmaDS, demonstrated higher recall. As for other approaches employed, they generally exhibited low performance.

For evaluating the performance of different DTU tools, we utilized precision, recall and F1 score (Table 2). For transcript-level analysis, we recommend using paired-end sequencing, as DTU analysis with single-end data captures less transcript information. When working with fewer than four replicates in paired-end data, we suggest using RSEM or Salmon as the quantification tool. Both tools perform similarly, but Salmon offers better runtime efficiency. If recall is a top priority, consider LimmaDS. If there are fewer replicates, DEXSeq could be a better choice. For more replicates, consider edgeR. When single-end sequencing is the only option, we also provided a guideline. For prioritizing recall, DSGseq can be employed. If there are fewer replicates, edgeR could be a better choice. For more replicates, consider DEXSeq. In addition, all tools’ performance is affected by the fold change and the number of transcripts. As the fold changes and the number of transcripts increases, recall of the tools generally increases. Most prominently, the precision of the tools decreases due to the greater number of S1 of DTU events found. S1 events are likely to be regulated by transcription factors rather than splicing. In our analysis, the tool with higher recall in S2 and S3 events and lower recall in S1 events is LimmaDS.

In our time series case study, we utilized both pairwise comparison tools that accommodate time series data and dedicated time series tools on a single-end time series transcriptome dataset from SARS-Cov-2 infected human cells. Interestingly, each tool identified different genes and few overlaps were observed. We observed that the pairwise comparison tools identified a substantial number of genes, exceeding 3000, whereas the time series tools reported fewer genes. Among the time series tools, only 48 genes were commonly identified. The Spycone method predominantly highlighted switching transcripts with high expression levels by calculating the event importance metrics. While TSIS found a lot of low expressed transcripts based on previous findings [37]. Furthermore, while Spycone focuses on detecting IS events, our simulation study revealed that both LimmaDS and DEXSeq do not differentiate between DTU scenarios. These observations suggest the importance of distinguishing S1 events, as doing so can yield unique insights. Even though the time series data is derived from single-end sequencing, it is shown that there are quantitative and qualitative differences while applying pairwise comparison tools and time series tools.

Greater effort is required to differentiate between DTU events scenarios, especially since these events are often mixed with varying degrees of change in real-world situations. In future work, we could leverage additional metrics from Spycone and apply them as filtering criteria to the results generated by pairwise comparison tools.

Exploring differential transcript usage (DTU) in single-cell data presents a fascinating avenue of study. Analyzing single cells offers a deeper understanding of transcript usage heterogeneity across various cell types. This can potentially reveal alternative splicing patterns that contribute to the emergence of distinct cell types. Several tools, such as DTUrtle and satuRn, have been developed specifically for detecting DTU in single-cell data, alongside with bulk RNA-seq. Our analysis indicates that while satuRn boasts a higher recall, its precision is marginally lower than that of DTUrtle. For datasets with a larger number of cells (>250 per cell type), DTUrtle is recommended for those prioritizing precision. Conversely, satuRn is the preferred option for datasets with fewer cells (<250 per cell type) and where recall is prioritized.

However, it’s worth noting that our analysis was based solely on a simulated dataset with an equal number of cells across two cell types. This balanced distribution does not always reflect real-world scenarios. Future analyses could benefit from simulating unbalanced datasets. Additionally, single-cell datasets often comprise more than just two cell types. As such, tools capable of comparing multiple cell types are more desirable. For instance, Acorde is designed to pinpoint co-DTU across several cell types [44]. Exploring DTU variations across pseudo-time within single-cell data presents a compelling direction for future research. However, single-cell transcript analysis faces technical challenges in obtaining accurate transcript counts. Ongoing developments in single-cell transcript analysis technologies suggest a promising future for understanding alternative splicing at the single-cell level [45,46].

**Table 2.** Recommended workflow for bulk single-end and paired-end data in DTU analysis. This workflow is for counts derived from different transcript quantification methods.

| Type of sequencing | Number of replicates | Quantification | Priority | Recommended tools |
| --- | --- | --- | --- | --- |
| paired-end | >8 | kallisto | precision | DEXSeq |
|  |  |  | recall | LimmaDS |
|  | <8 |  | precision | DEXSeq |
|  |  |  | recall | DEXSeq |
| paired-end | >8 | RSEM | precision | satuRn |
|  |  |  | recall | LimmaDS |
|  | <8 |  | precision | satuRn |
|  |  |  | recall | LimmaDS |
| paired-end | >8 | Salmon | precision | satuRn |
|  |  |  | recall | LimmaDS |
|  | <8 |  | precision | satuRn |
|  |  |  | recall | LimmaDS |
| single-end | >8 | Kallisto | precision | DEXSeq |
|  |  |  | recall | DEXSeq |
|  | <8 |  | precision | DEXSeq |
|  |  |  | recall | DEXSeq |
| single-end | >8 | RSEM | precision | seqGSEA |
|  |  |  | recall | seqGSEA |
|  | <8 |  | precision | LimmaDS |
|  |  |  | recall | LimmaDS |
| single-end | >8 | Salmon | precision | seqGSEA |
|  |  |  | recall | seqGSEA |
|  | <8 |  | precision | LimmaDS |
|  |  |  | recall | LimmaDS |

## Conclusion

In this comprehensive benchmark study, we have rigorously evaluated various transcriptomics datasets under both static and dynamic conditions, utilizing a blend of simulated and real data. We observed a general trend where tools using generalized linear models or Dirichlet-multinomial models showed superior precision, while LimmaDS, which employs a linear model, demonstrated higher recall. This suggests that the choice of DTU tools should be tailored to the specific needs of the study, considering factors like the number of replicates, sequencing method (single-end or paired-end), and the prioritization of precision or recall. Our analysis of time series data revealed interesting insights into DTU. Tools like Spycone, designed for time series DTU detection, showed differences in functional outcome.

Looking ahead, there is a clear need for further differentiation between DTU events in complex real-world scenarios. Additionally, exploring DTU in single-cell data remains a promising avenue, albeit with technical challenges in obtaining accurate transcript counts. As single-cell transcript analysis technologies continue to evolve, they hold significant promise for advancing our understanding of alternative splicing at the single-cell level. Future research could benefit from simulating more diverse and unbalanced datasets, as well as focusing on tools capable of comparing multiple cell types and analyzing DTU variations across pseudo-time.

## Key points

- We performed an analysis involving a comprehensive benchmark study that used both simulated data and real transcriptomics datasets. It considered both static and dynamic conditions. We included the 12 DTU tools for static conditions. Our study suggests that tools using generalized linear models produce better precision and with linear models produce better recall.
- Based on the stratifications of the DTU genes, recall of most tools are positively affected by the fold changes and number of transcripts. Our results show that LimmaDS is better in detecting S2 and S3 events scenarios.
- We provided guidelines for performing DTU analysis for different sequencing types, considering the number of replicates. LimmaDS, edgeR and DEXSeq are better for paired-end sequencing. DSGseq and DEXSeq are better for single-end sequencing.
- We provided evidence that Spycone can detect IS that has a different biological interpretation to the condition of interest.
- For datasets containing more than 250 cells per cell type, DTUrtle is the suggested choice for those valuing precision. On the other hand, for datasets with less than 250 cells per cell type, satuRn is recommended when recall is of greater importance.

## Author contributions

C.T.L. planned and carried out the analysis. T.D. performed the single-cell analysis. M.H. and L.W. preprocessed the SARS-CoV2 datasets. C.T.L., M.L., M.H., O.T., and J.B. wrote and reviewed the manuscript.

