## Supplementary material for "Comprehensive benchmark of differential transcript usage analysis for static and dynamic conditions": Figure S2

Supplementary figures

**Effect of different transcript quantification method**

The results from isoform quantifications tools Salmon, Kallisto, and RSEM are compared by RMSE, Spearman, and Pearson correlation coefficients. The following formula calculates the Root mean square error (RMSE):

$RMSE = \sqrt{\sum(E{i - T{i){}^{2}/ n}}}$	(3)

where E_i_ is the estimated count of transcript i, T_i_ is the groundtruth of transcript i and n is the number of transcripts.

The following formula calculates the Spearman correlation:

$Spearman = \frac{6\sum(d{i){}^{2}}}{n(n{{}^{2}}-1)}$	(4)

where d_i_ is the difference of the rank between ground truth and estimated counts. The sum of the differences is then rescale to the final correlation coefficient based on the number of transcripts n.

The following formula calculates the Pearson correlation:

$r = \frac{cov(E{}_{i}, T{}_{i})}{\sigma{}_{E}, \sigma{}_{T}}$	(5)

where E_i_ is the estimated count of transcript i and T_i_ is the groundtruth of transcript i. Standard deviation is calculated for both estimated counts and groundtruth.

RSEM, Salmon and kallisto are among commonly used methods for transcript quantification. Salmon and kallisto perform fast pseudo-alignment using k-mers, whereas RSEM requires already aligned reads as input. We evaluated the transcript quantification of the simulated datasets by comparing the counts estimated by the methods to the ground truth. The ground truth is the modified transcript counts that were passed to the rsem-simulate-reads function. To evaluate, we calculated root mean square error (RMSE) between the simulated ground truth and the estimated counts by the quantification methods (Figure. S1). The RMSEs in paired-end data are around ten-fold lower than in single-end data, emphasizing the considerable quality gain of paired-end sequencing. Moreover, kallisto counts have two times higher RMSE than those reported by RSEM and Salmon. We also calculated the Pearson and Spearman correlation against the simulated ground truth (Figure. S1). Pearson correlation assesses the linearity of the counts, while Spearman is a rank-based measure. For paired-end data, all three quantification methods perform similarly in terms of Pearson correlation. Salmon is slightly better in terms of Spearman correlation in paired-end data. In single-end data, kallisto had a Pearson correlation of 0.2 but a 0.75 Spearman correlation. This result suggested that kallisto can overestimate or underestimate counts, while keeping the ranking of the genes intact. For RSEM and Salmon, RMSE and Spearman correlation suggest opposite results. RSEM is better in terms of Spearman correlation, however, Salmon is better in terms of RMSE.

**Differential transcript usage detection tools**

Exon-centric tools focus on counting bins of reads overlapping with the exons/junctions. DEXSeq (v1.42.0) and JunctionSeq (v1.5.4) both use a generalized linear model (GLM) to test for differential usage of exons and splice junctions, respectively. Transcript-centric tools make use of transcript count data. For DEXSeq, conditions are compared using the design formula *~sample+exon+group:exon*. Since DEXSeq was initially designed for modeling exon counts, it can be adapted for transcript counts by supplying a transcript count matrix. To filter significant genes, per gene Q values (equivalent to adjusted p-value) are calculated. To evaluate JunctionSeq, we used the resulting gene-wise adjusted p-value for further evaluation. DSGSeq (v0.1.0) calculates negative binomial statistics by comparing exon-counts between two conditions. seqGSEA (v1.36.0) combines the DSGseq and DESeq methods by generating a normalized metric of DS and DE score using a rank-based strategy. seqGSEA uses exon counts from HTSeq and requires an exon annotation derived from RefSeq or Ensembl annotation reference. SeqGSEA outputs a p-value by permuting the DS score. Genes that have an adjusted p-value lower than 0.05 are considered DTU genes for further evaluation.

For transcript-centric tools, DRIMSeq (v1.24.0) uses a dirichlet-multinomial model to test for differential distribution of transcript abundances. The resulting adjusted p-values are used for DTU gene filtering. DTUrtle (v1.0.2) extends DRIM-Seq with extra post-hoc filtering steps prior to using StageR. StageR corrects false-discovery rate on gene-level instead of hypothesis level. As a result, an overall false-discovery rate was obtained per gene. Iso-KTSP (v1.0.3) tests for switching transcript pairs that separate different conditions by using a classifier, e.g. tumor vs normal samples. Each pair of transcripts in a gene is scored based on the frequency of how often one transcript has higher expression than other in one condition. Based on this score, the classifier determines if the transcript pair belongs to the ‘tumor’ or ‘normal’ group. For each transcript pair, an information gain value is calculated to indicate the predictive power given the samples that are correctly predicted as a tumor/normal sample. satuRn uses a quasi-binomial generalized linear model to model the transcript counts of a gene. The resulting adjusted p-values are used for DTU gene filtering. Cufflinks/cuffdiff (v2.2.1) is a suite of tools that perform de novo transcript alignment and differential expression analysis at both gene and transcript levels. Cuffdiff uses a poisson model that models the variability of biological replicates in terms of fragment counts in a transcript. The selected tools are summarized in table 2.

For all the tools that provide adjusted p-values as results, genes with values lower than 0.05 are considered significant. Iso-KTSP and DSGseq, which do not provide p-values, use 0.5 and 5 respectively as cutoffs chosen based on the recommendations of the authors.

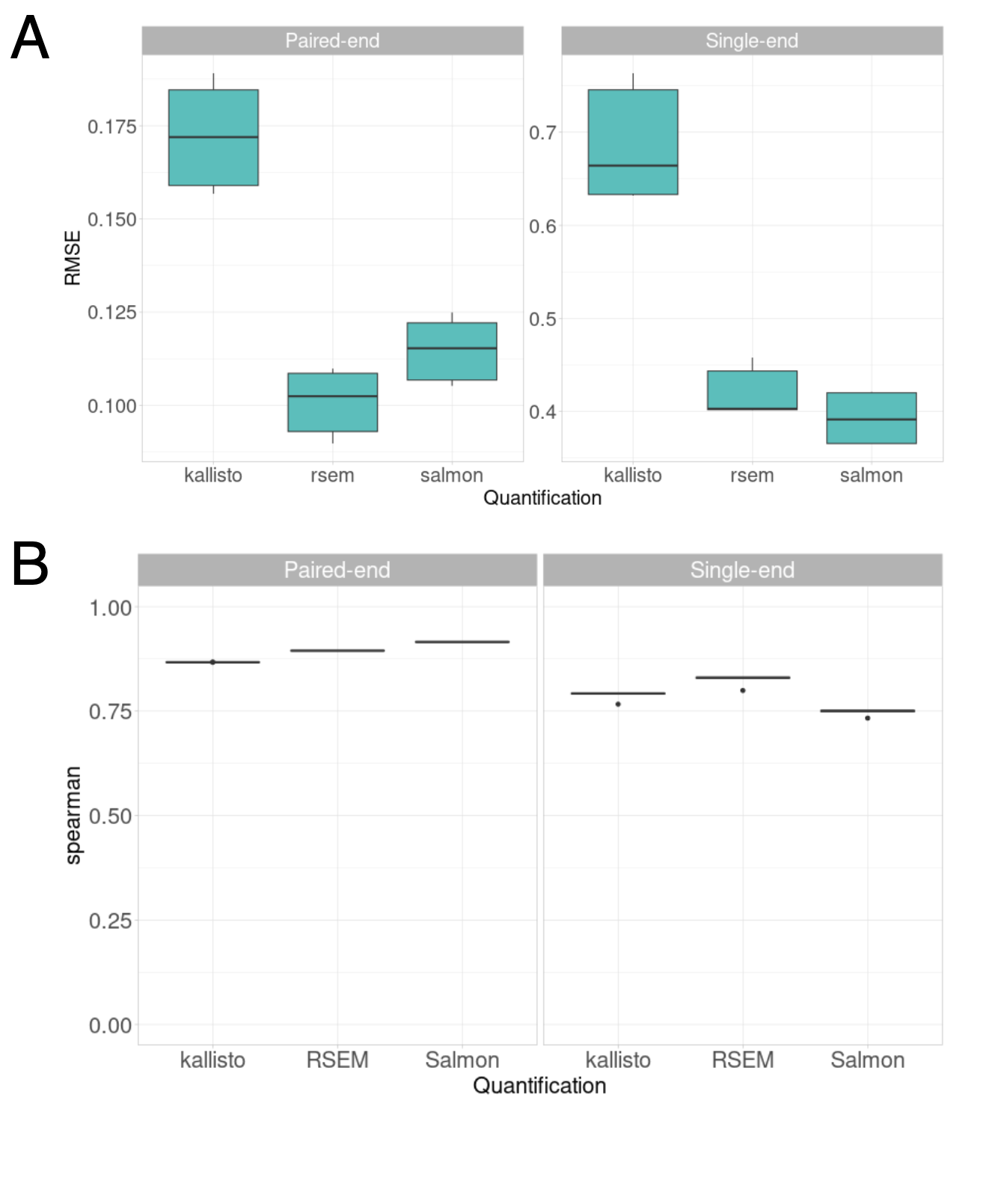

Figure S1. A) Boxplots showing the RMSE of the estimated counts from all three transcript quantification tools in both paired-end and single-end data. B) Spearman correlation between estimated transcript count and groundtruth.

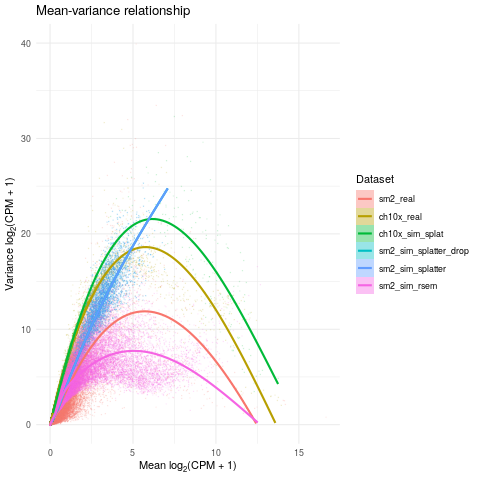

Figure S2. Comparison on simulated data by RSEM and splatter in Smart-seq2 data and Chromium 10x data. a

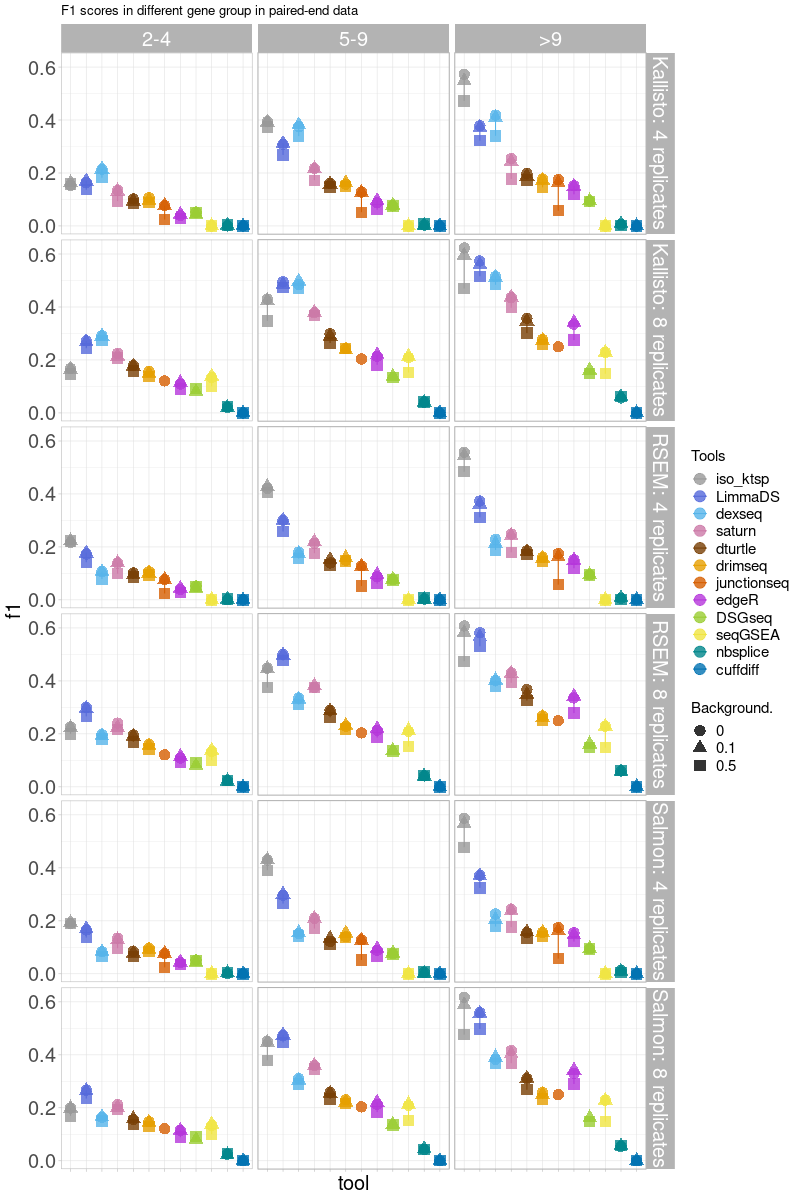

Figure S3. F1 scores are stratified by the number of isoforms of all combinations of quantification tools and DTU methods in paired-end data with 4 replicates (top row) and 8 replicates (bottom row).

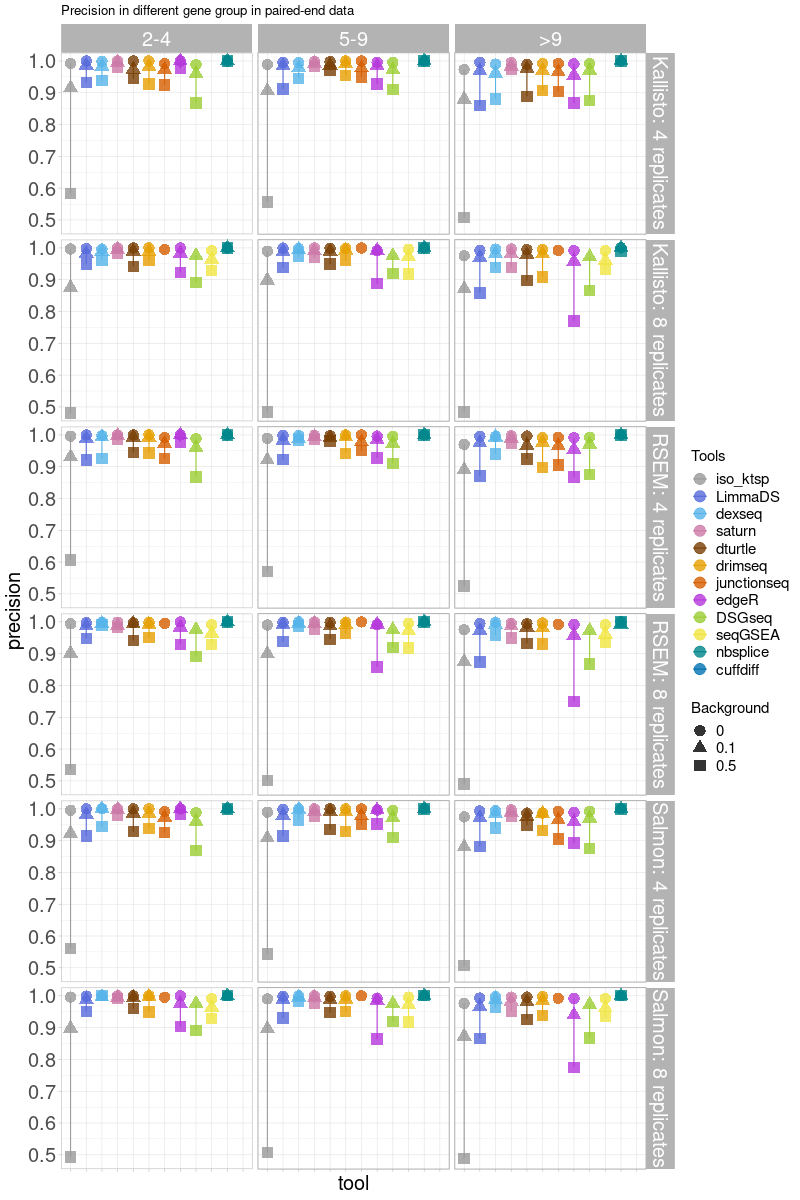

Figure S4. Precision stratified by the number of isoforms of all combinations of quantification tools and DTU methods in paired-end data with 4 replicates (top row) and 8 replicates (bottom row).

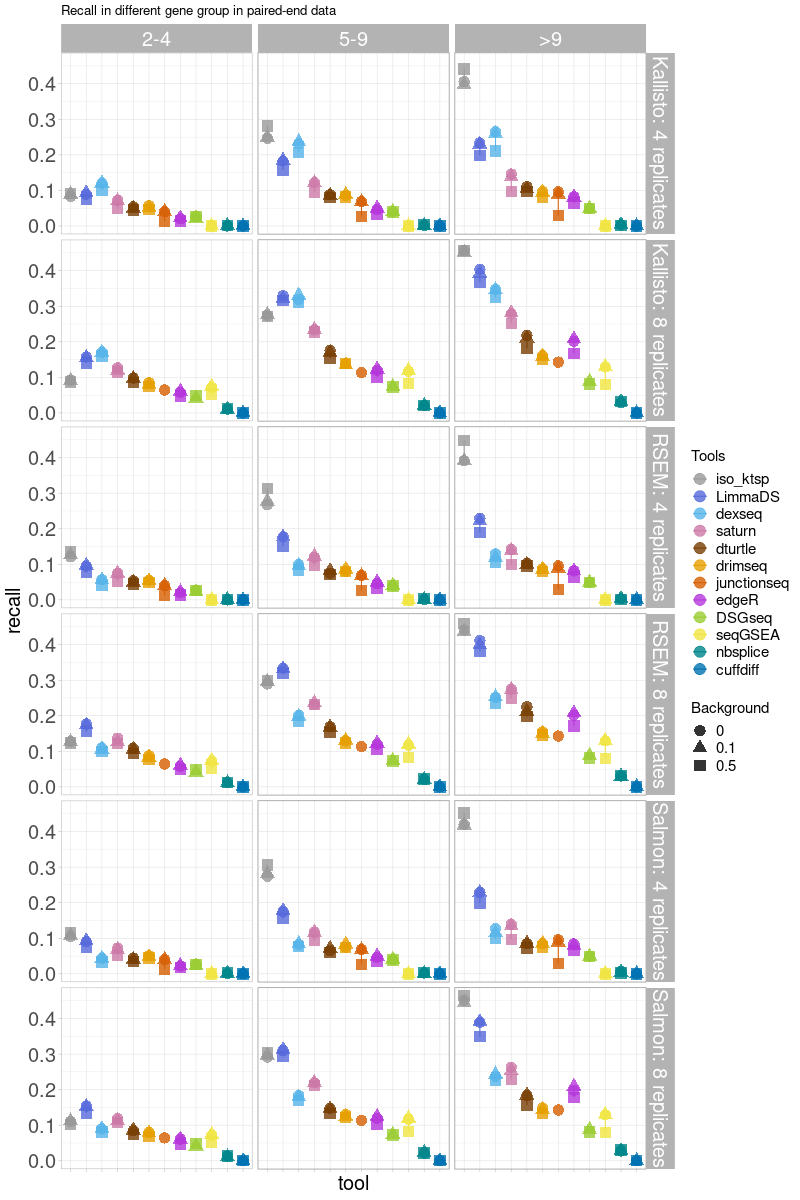

Figure S5. Recall stratified by the number of isoforms of all combinations of quantification tools and DTU methods in paired-end data with 4 replicates (top row) and 8 replicates (bottom row).

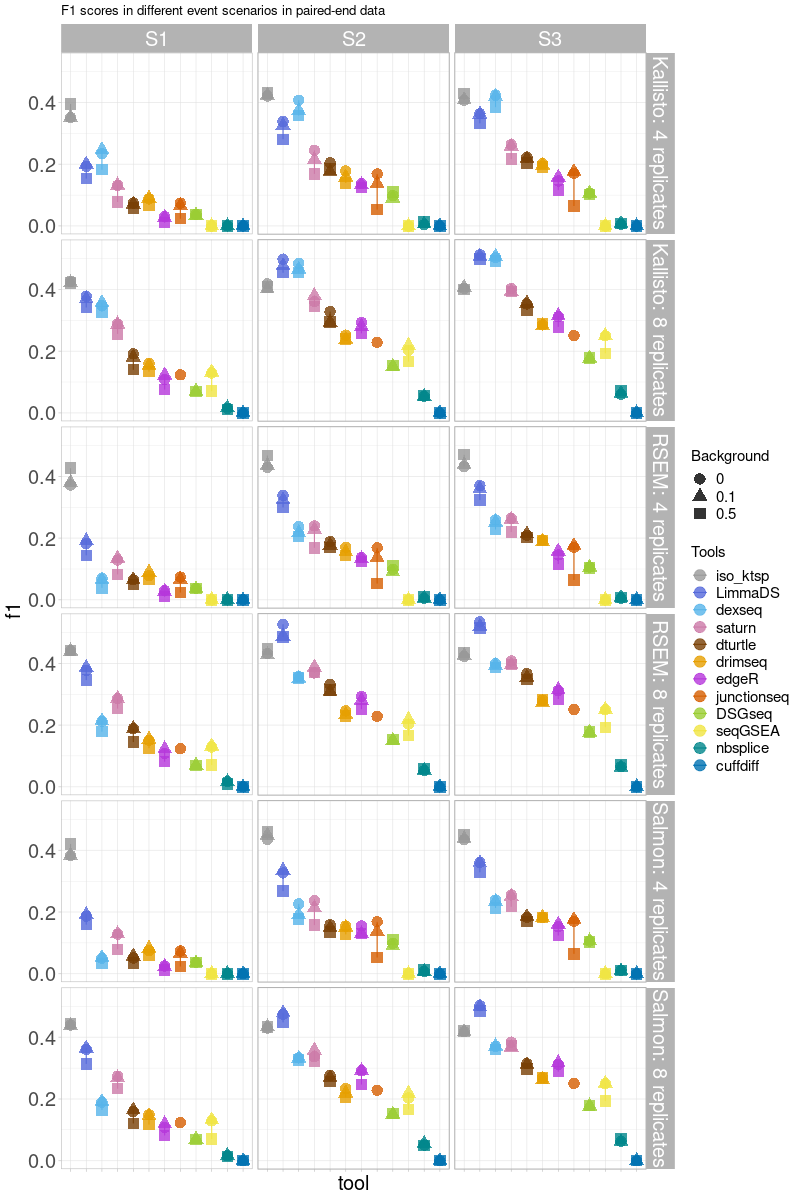

Figure S6. F1 scores are stratified by **fold change** of all combinations of quantification tools and DTU methods in **paired-end data** with 4 replicates and 8 replicates.

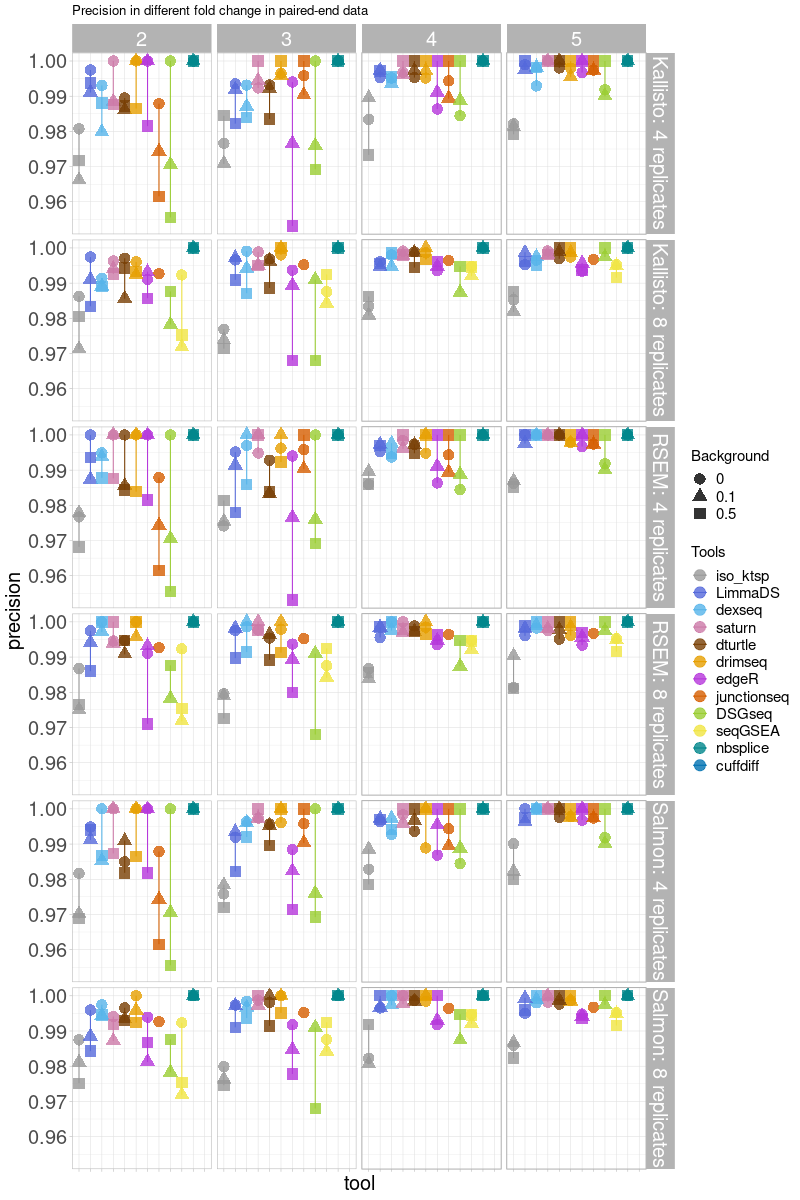

Figure S7. Precision stratified by fold change of all combinations of quantification tools and DTU methods in paired-end data with 4 replicates and 8 replicates.

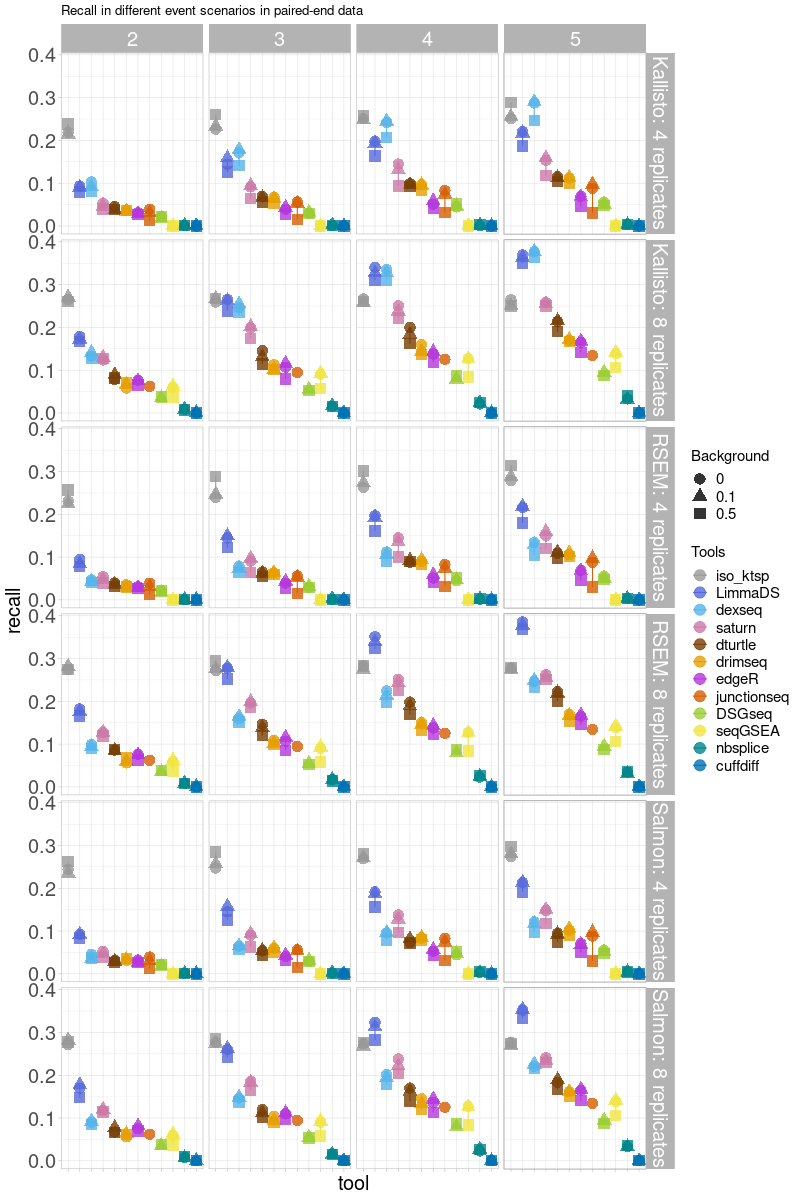

Figure S8. Recall stratified by fold change of all combinations of quantification tools and DTU methods in paired-end data with 4 replicates and 8 replicates.

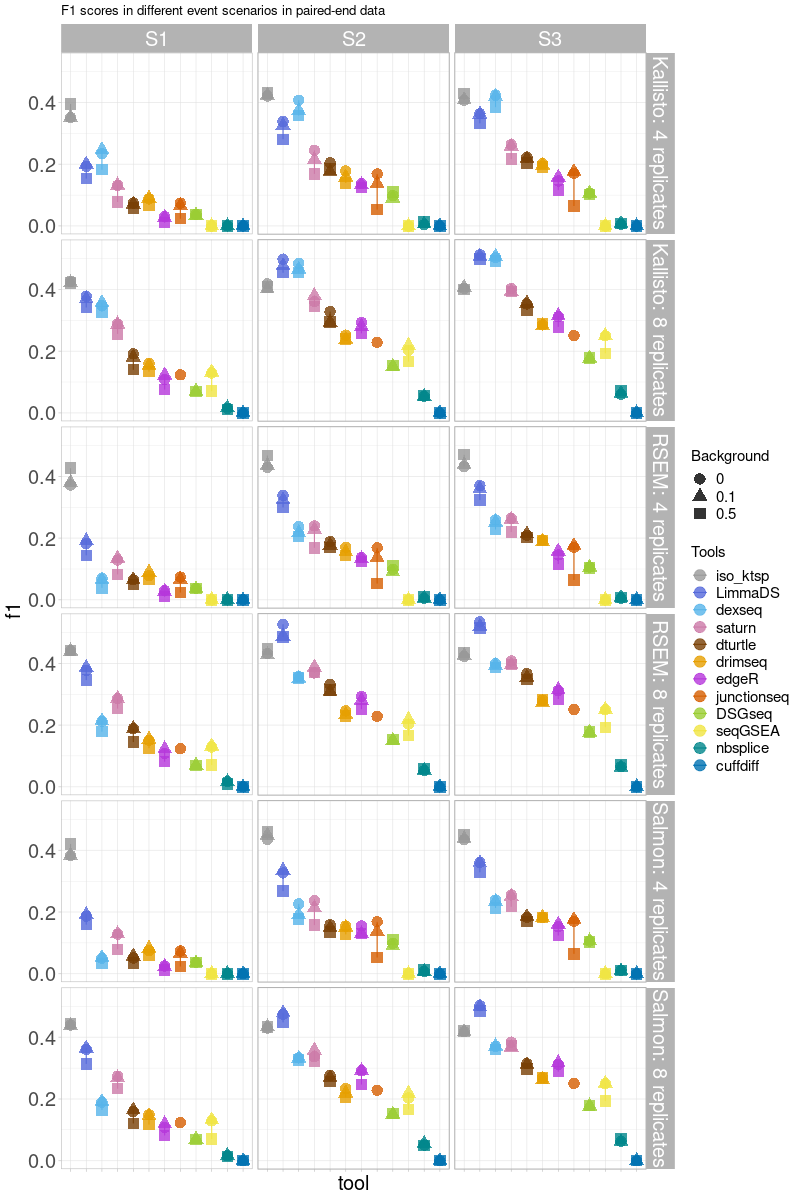

Figure S9. F1 scores stratified by different event scenarios of all combinations of quantification tools and DTU methods in paired-end data with 4 replicates and 8 replicates.

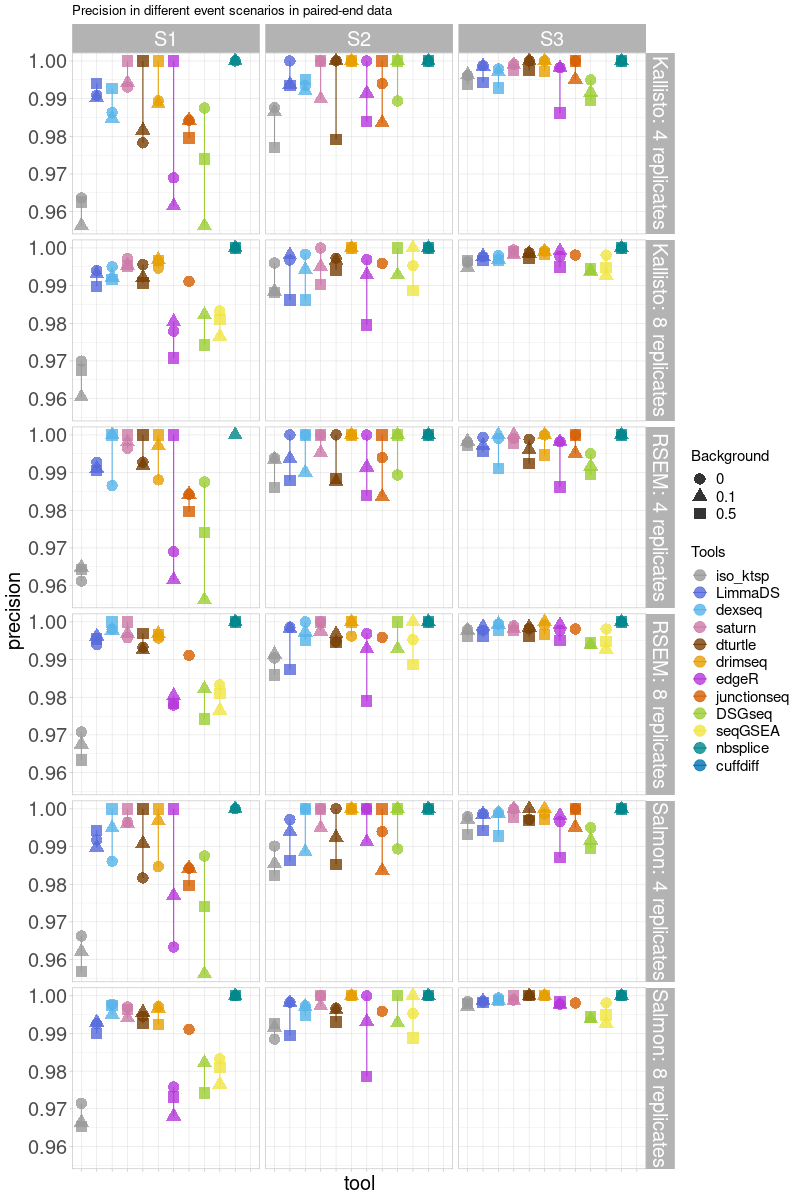

Figure S10. Precision stratified by different event scnenarios of all combinations of quantification tools and DTU methods in paired-end data with 4 replicates and 8 replicates.

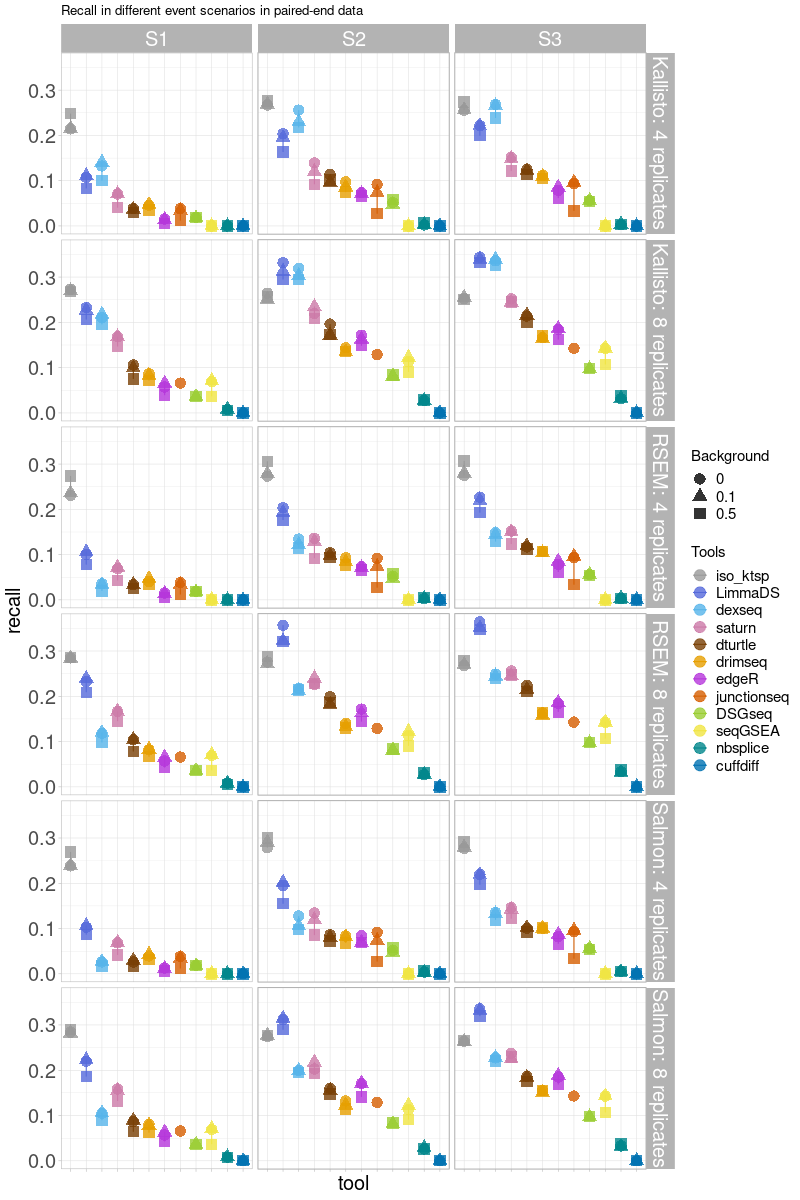

Figure S11. Recall stratified by different event scenarios of all combinations of quantification tools and DTU methods in paired-end data with 4 replicates and 8 replicates.

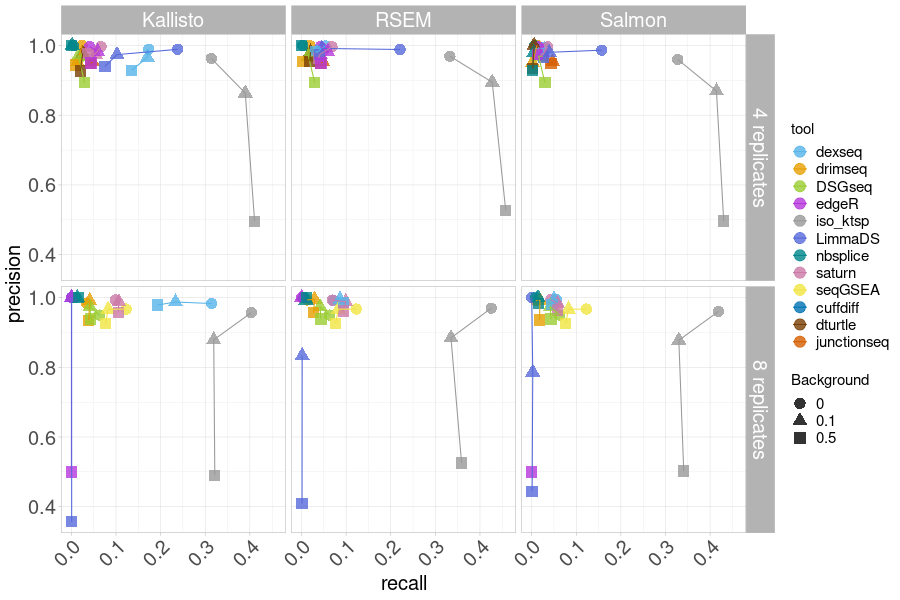

Figure S12. Precision and recall plot of all combinations of quantification tools and DTU methods in **single-end data** with 4 replicates (left) and 8 replicates (right).

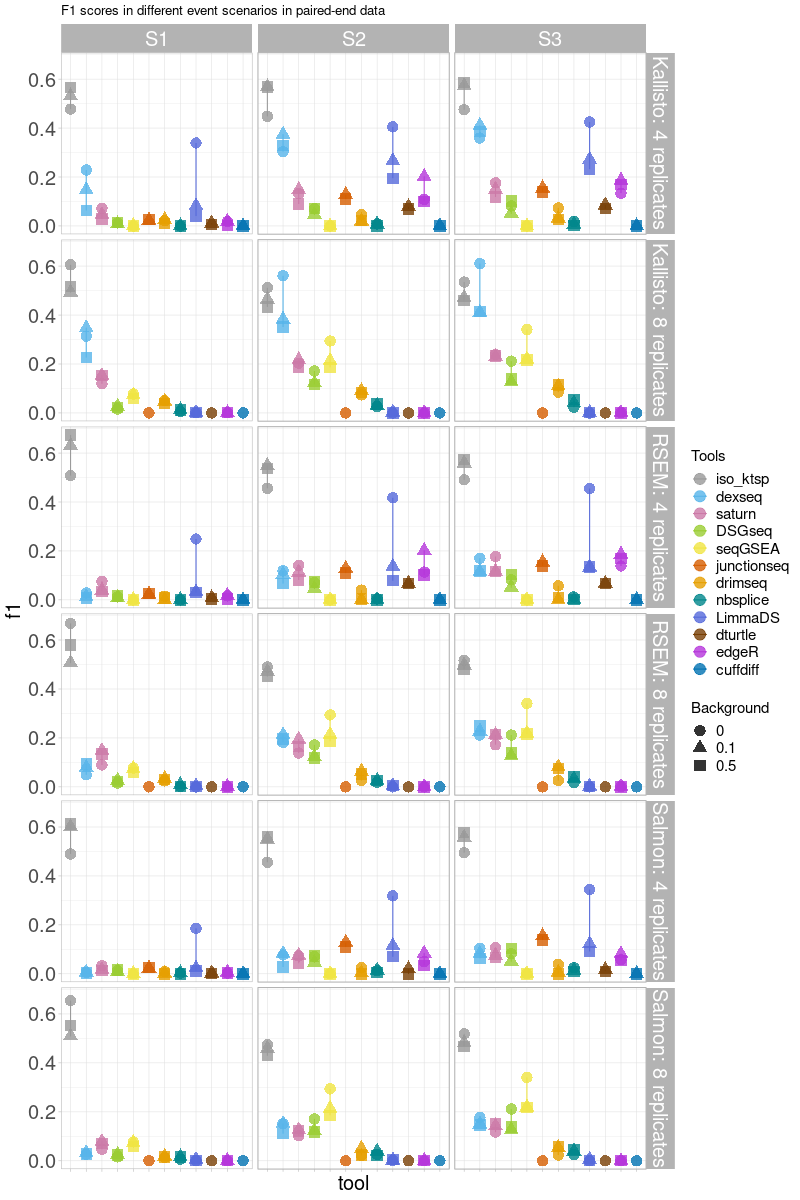

Figure S13. F1 scores plot stratified by different event scenarios of all combinations of quantification tools and DTU methods in **single-end data** with 4 replicates (top row) and 8 replicates (bottom row).

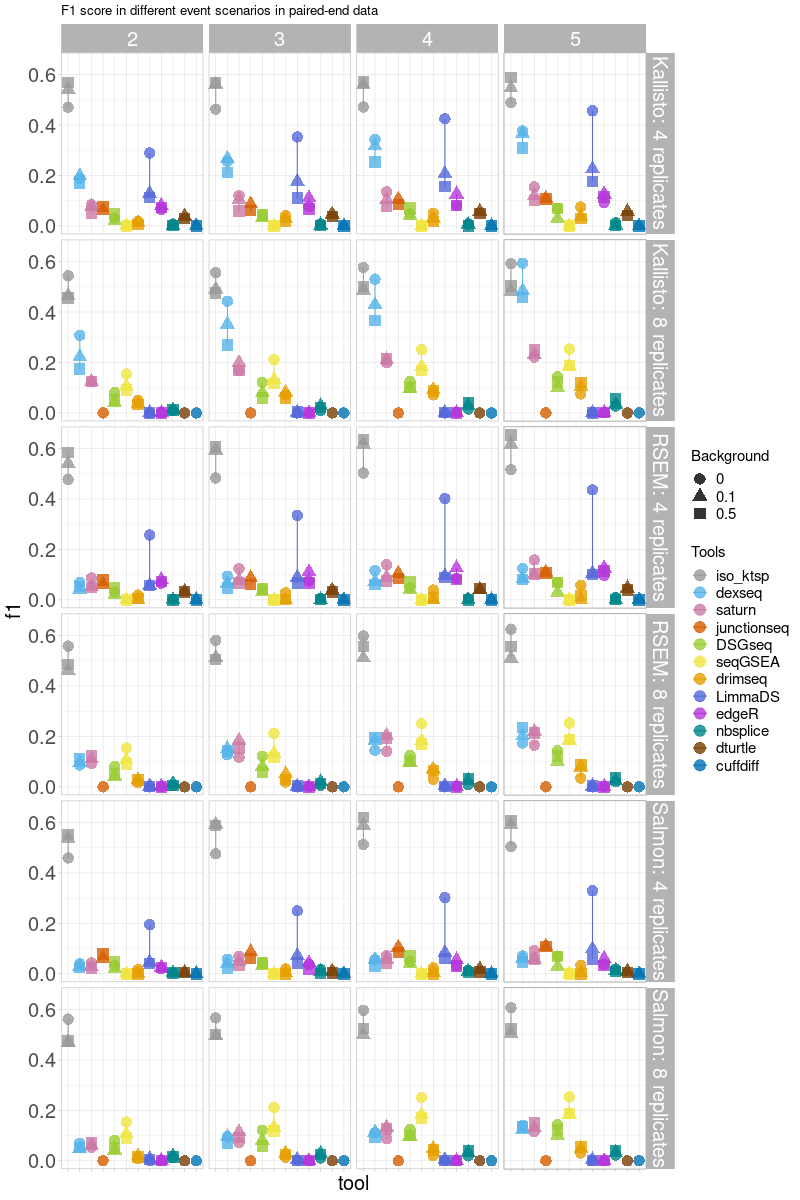

Figure S14. F1 scores plot stratified by fold change of all combinations of quantification tools and DTU methods in **single-end data** with 4 replicates (top row) and 8 replicates (bottom row).

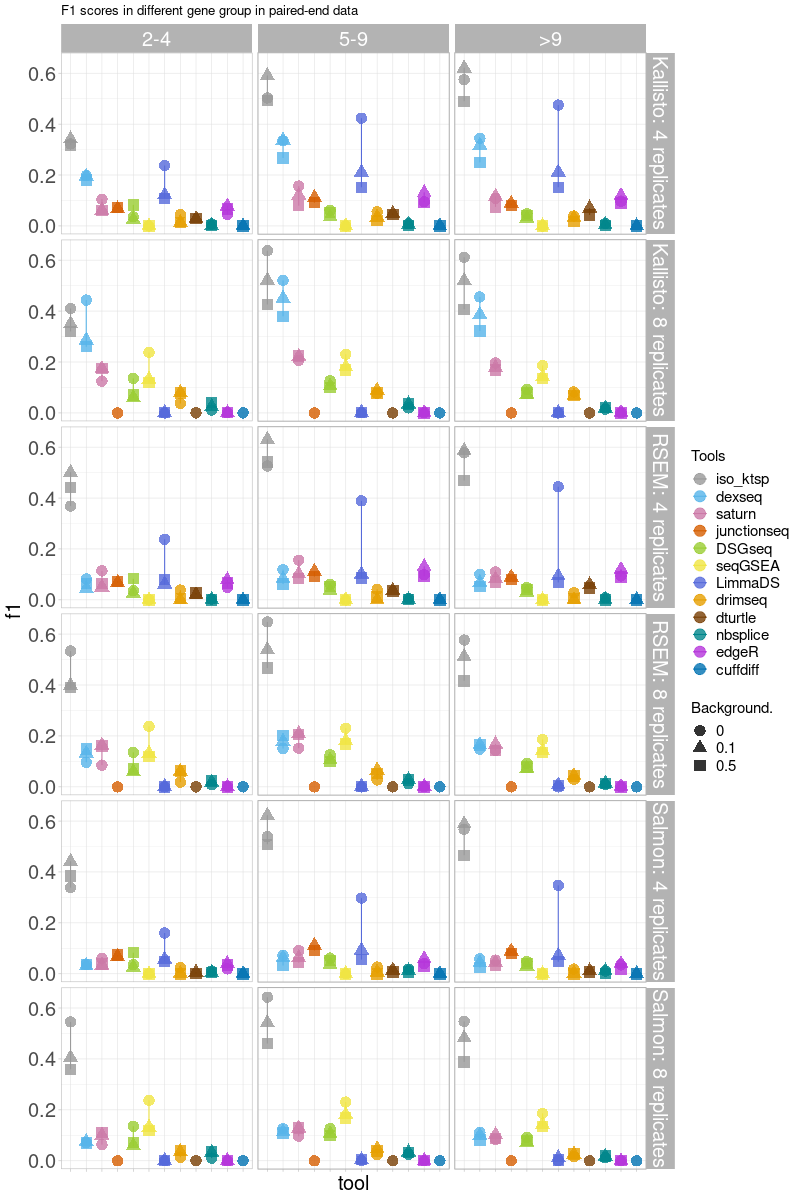

Figure S15. F1 scores plot stratified by number of transcripts of all combinations of quantification tools and DTU methods in **single-end data** with 4 replicates (top row) and 8 replicates (bottom row).

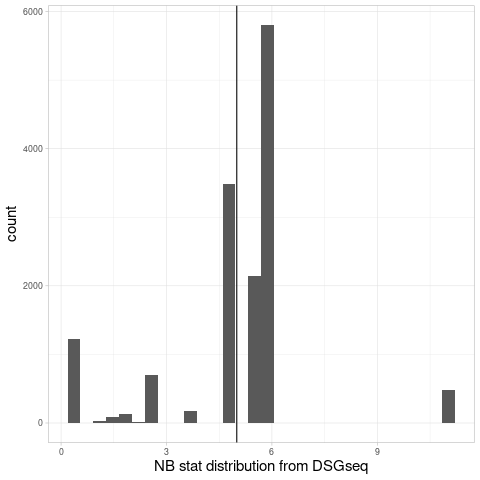

Figure S16. The distribution of NB statistics from DSGseq derived from the prostate cancer dataset. The threshold of 5 is chosen based on this plot.

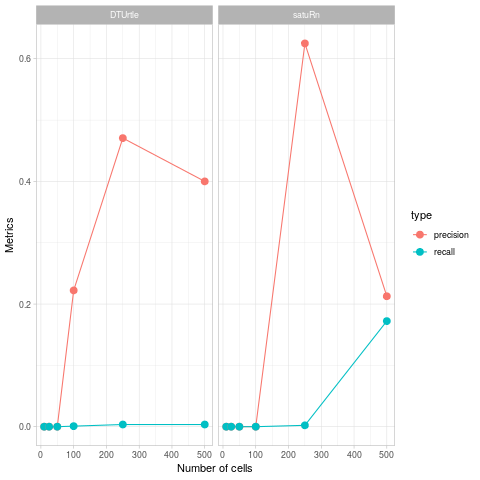

Figure S17. Pseudo-bulk analysis of the simulated single-cell dataset.

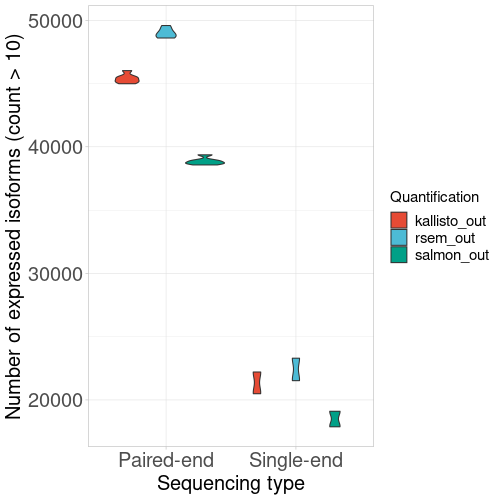

Figure S18. The number of transcripts detected in the quantification methods based on sequencing type.

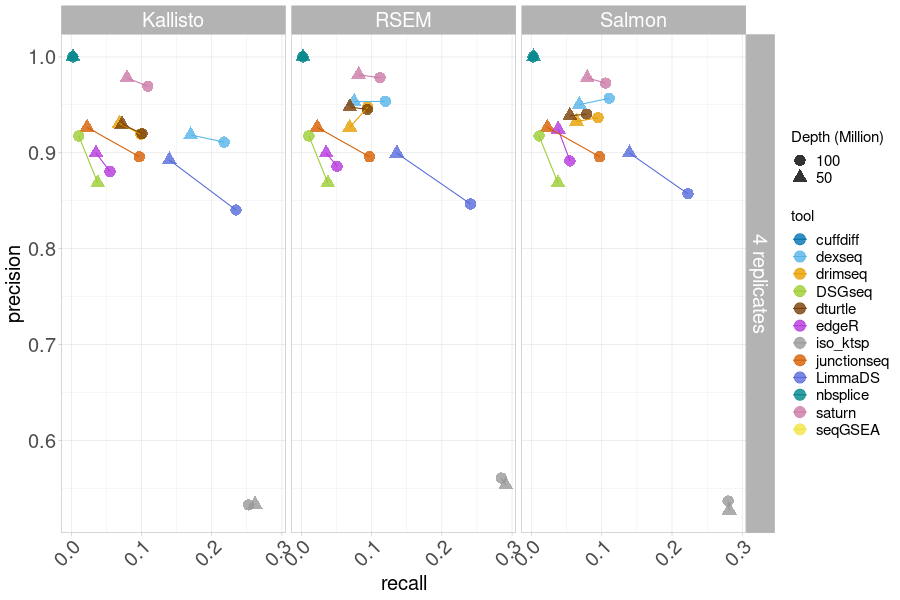
Figure S19. Precision and recall plot comparing **50M and 100M sequencing depth** of all combinations of quantification tools and DTU methods in **paired-end data** with 4 replicates at 0.5 background level.
